# Vocal biomarkers of aging and Parkinson’s disease in a songbird

**DOI:** 10.64898/2026.07.26.740830

**Authors:** Lucas M. Dal’Ava, Plinio A. Barbosa, Julie E. Miller

**Author notes:** Corresponding Author: L. M. Dal’Ava.

## Abstract

**Purpose:** Similar vocal impairments occur in normative aging and Parkinson’s disease (PD), making differential diagnosis challenging in early stages. The brain’s role in age and PD-related vocal dysfunction is unclear. This motivated our use of the zebra finch songbird model. We applied human acoustic analysis to evaluate aging and parkinsonian-like birdsong for shared and distinct features.

**Method:** Two open-access datasets of zebra finch song were analyzed using an expanded set of parameters that, in the case of humans, can detect subtle changes in voice. The aging dataset included younger, middle aged, and older birds. The PD dataset included birds receiving viral injections into Area X to induce alpha-synuclein overexpression (ASYN) or control treatment, recorded pre- and post-injection. Songs were automatically segmented into motifs using cross-correlation. Acoustic measures were extracted from harmonic, noisy, and mixed syllables.

**Results:** Middle aged birds differed from younger and older birds, showing non-linear trajectories in fundamental frequency (*f*_o_) measures and age-related increases in spectral slope, spectral emphasis, and intensity variability, whereas temporal parameters remained stable. In PD, pre-post changes occurred in both ASYN and controls, with ASYN birds exhibiting larger effects for *f*_o_ variability, decreasing frequency modulation, and vocal intensity measures.

**Conclusions:** Vocal aging in zebra finches was characterized by non-linear trajectories and prominent spectral and intensity-related changes. In a gene-driven PD state there were greater alterations in *f*_o_ variability and intensity regulation, features that may distinguish aging- from ASYN-related vocal variability. These findings support the zebra finch as a translational model for vocal biomarkers research.

## Introduction

The aging population is one of the most significant demographic transformations worldwide. Driven by declining fertility rates and increasing life expectancy, the number of individuals aged 65 years and older is projected to more than double in coming decades, reshaping healthcare demands and social systems (UNDESA, 2023).

Voice disorders are prevalent among older adults. Approximately 18-20% of older adults experience clinically significant voice disorders, with prevalence increasing with age (Wang *et al*., 2023). Age-related vocal changes, collectively referred to as presbyphonia, include increased breathiness, roughness, vocal weakness, tremor, reduced intensity, and shifts in fundamental frequency (*f*_o_) (Galluzzi & Garavello, 2018; Beton *et al*., 2022). These changes co-occur with declines in speech motor control, including reduced articulatory precision, slower speech rate, and increased temporal variability, contributing to decreased speech intelligibility.

Communication impairments in aging are associated with reduced quality of life, social withdrawal, and adverse mental health outcomes (Palmer *et al*., 2019). Despite these consequences, most research has focused on peripheral laryngeal mechanisms, with comparatively less attention given to central neural mechanisms underlying vocal motor control during aging (Vasquez *et al*., 2026).

In humans, vocal production depends on the coordinated functioning of multiple physiological systems, including respiratory, muscular, sensory, and neural systems. Phonation, in particular, requires precise coordination between respiration and the laryngeal musculature (Veit *et al*., 2011; Park *et al*., 2024). Aging affects these systems and is associated with declines in respiratory and laryngeal muscle performance across humans and several animal species, including baboons and rats (Mardini *et al*., 1987; McMullen & Andrade, 2006; Lenell *et al*., 2018; Peres *et al*., 2024). Importantly, age-related vocal changes do not necessarily follow a uniform trajectory; non-linear changes in *f*_o_, including U-shaped trajectories, have been reported across adulthood in humans (Stathopoulos *et al*., 2011; Valente *et al*., 2024).

In addition to normative aging, neurodegenerative conditions may exacerbate age-related vocal changes in humans. Parkinson’s disease (PD), characterized by alpha-synuclein aggregation and dopaminergic degeneration in the basal ganglia, provides a framework for understanding central neural dysfunction in vocal behavior. PD impairs motor timing, coordination, and sensorimotor integration, manifesting in speech and voice production. PD-related speech changes are also influenced by respiratory and laryngeal alterations affecting airflow, intensity, and phonatory stability (Sapir, 2014; Darling-White *et al*., 2022). Vocal deficits are highly prevalent in PD and often emerge early, highlighting the sensitivity of vocal production to basal ganglia dysfunction (Sapir, 2014; Polychronis *et al*., 2026).

While age- and PD-related vocal changes have been extensively studied in humans, fewer studies have examined these processes in non-human species. Animal models allow disentangling normative aging from disease-specific neural degeneration under controlled conditions, which is particularly valuable for vocal behaviors that rely on conserved principles of motor sequencing, timing, and basal ganglia modulation.

Songbirds provide a compelling model system for investigating vocal aging and neurodegeneration as vocal learning and production rely on a well-characterized network of cortical and basal ganglia nuclei. In zebra finches (*Taeniopygia guttata*), song production is restricted to males, and these individuals continue to produce learned songs throughout adulthood and advanced age, allowing lifespan-related changes in vocal motor behavior to be examined without behavioral cessation (Brainard & Doupe, 2013; Cooper *et al*., 2012; Badwal *et al*., 2020; Berg *et al*., 2020). Consistent with this, aging in male zebra finches is associated with changes in *f*_o_, amplitude, spectral structure, and vocal variability, providing a sensitive model for detecting age-related changes in vocal motor output (Cooper *et al*., 2012; Badwal e*t al*., 2020; Higgins *et al*., 2025; Gordon *et al*., 2025).

Beyond age-related changes, zebra finches have emerged as a robust translational model for investigating PD-related vocal dysfunction. Overexpression of the human alpha-synuclein (h*SNCA*) gene in cortico-basal ganglia circuits induces significant changes in song timing, spectral organization, and acoustic variability, mirroring key features of hypokinetic dysarthria in PD (Medina *et al*., 2022; Bjork *et al*., 2025).

The translational relevance of songbirds is further supported by shared organizational principles of vocal motor control across species. Songbirds possess cortical and basal ganglia circuits with functional and molecular parallels to human speech-related networks (Pfenning *et al*., 2014). In humans, direct cortical control over laryngeal motor neurons enables fine-grained modulation of *f*_o_ and intensity (Zhang *et al*., 2023), reinforcing comparative approaches for understanding vocal communication.

This study investigates how normative aging and PD-related pathology shape vocal motor control in zebra finches using a comparative framework grounded in human acoustic-prosodic analysis. Two previously published datasets were analyzed: one examining age-related vocal changes (Badwal *et al*., 2020) and another employing an alpha-synuclein overexpression model of PD-related vocal dysfunction (Medina *et al*., 2022). We automated birdsong segmentation and expanded the range of acoustic parameters, applying an extensive set of acoustic-prosodic measures commonly used in human voice research (Dal’Ava & Barbosa, 2024; Dal’Ava, 2025).

We hypothesized that (a) aging would be associated with measurable changes across multiple acoustic domains, particularly spectral, intensity, and *f*_o_-related measures, and (b) alpha-synuclein overexpression would produce additional alterations in temporal, spectral, and *f*_o_-related measures reflecting deficits in fine vocal motor control. By integrating human voice analysis methods with songbird vocal data, this study aims to identify overlapping and distinct acoustic signatures associated with aging and PD-related neural dysfunction, and further evaluate the translational relevance of the zebra finch model for vocal biomarker research both in humans and birds.

## Method

### Datasets

All recordings were obtained from two previously published open-access datasets collected from adult male zebra finches in the Miller colony at the University of Arizona between 2015 and 2020 (Badwal *et al*., 2020; Medina *et al*., 2022) (see Data Availability statement). These datasets were collected under author J.E. Miller’s approved protocol #13-489 by the Institutional Animal Care and Use Committee at the University of Arizona. In the aging dataset, birds were naïve and grouped by age based on their natural lifespan in the laboratory colony: Younger adults (241-294 days post-hatch, dph, n = 8), middle aged adults (627-898 dph, n = 8), and older adults (1147-1402 dph, n = 10) (Badwal *et al*., 2020). In the PD model dataset, adult male zebra finches (120-400 dph, n = 15) were assigned to an alpha-synuclein overexpression group (ASYN; n = 9) or a control group (GFP, n = 6), receiving bilateral viral injections into Area X, a song-dedicated basal ganglia nucleus, with either the human alpha-synuclein (h*SNCA*) gene or viral placebo (green fluorescent protein - GFP).

### Recording Parameters

All recordings of individual males were conducted in sound-attenuated chambers during two-hour morning sessions immediately following lights-on using Shure omnidirectional microphones at a sampling rate of 44.1 kHz and 24-bit resolution. Zebra finch songs were recorded under undirected (UD) conditions, i.e., in the absence of a female, a context known to elicit increased variability in acoustic parameters compared to directed song (Sober *et al*., 2008). Two hours of UD recordings were collected each morning under controlled light conditions using Song Analysis Pro (SAP) 2011 (Tchernichovski *et al*., 2000). While singing, birds typically remain stationary within their cages; previous observations indicate that positional changes introduce minimal intensity variation (approximately 1 dB). To ensure comparable intensity measurements across recording chambers, each chamber was acoustically calibrated using a 1,000-Hz reference tone. The intensity of the calibration tone was measured at the microphone using a sound level meter and subsequently quantified in Praat, allowing the derivation of chamber-specific calibration factors that were applied to all intensity measurements (Badwal *et al*., 2020). Recordings were automatically triggered when signal amplitude exceeded predefined thresholds above the ambient noise floor, and songs were segmented according to minimum syllable duration and maximum allowable inter-syllable silence parameters (Tchernichovski *et al*., 2000; Badwal *et al*., 2020; Medina *et al*., 2022).

### Acoustic Data Extraction and Segmentation

A motif was defined as a stereotyped sequence of syllables repeatedly produced by an individual bird during song. In zebra finches, a syllable is acoustically defined as a discrete sound unit separated from adjacent units by a local minimum in the amplitude envelope (i.e., a silent pause or a marked drop in acoustic energy); spectrally, syllables can exhibit harmonic structure, broadband noise, or a mixture of both (Badwal *et al*., 2019) (see Figure 1).

**Figure 1.**
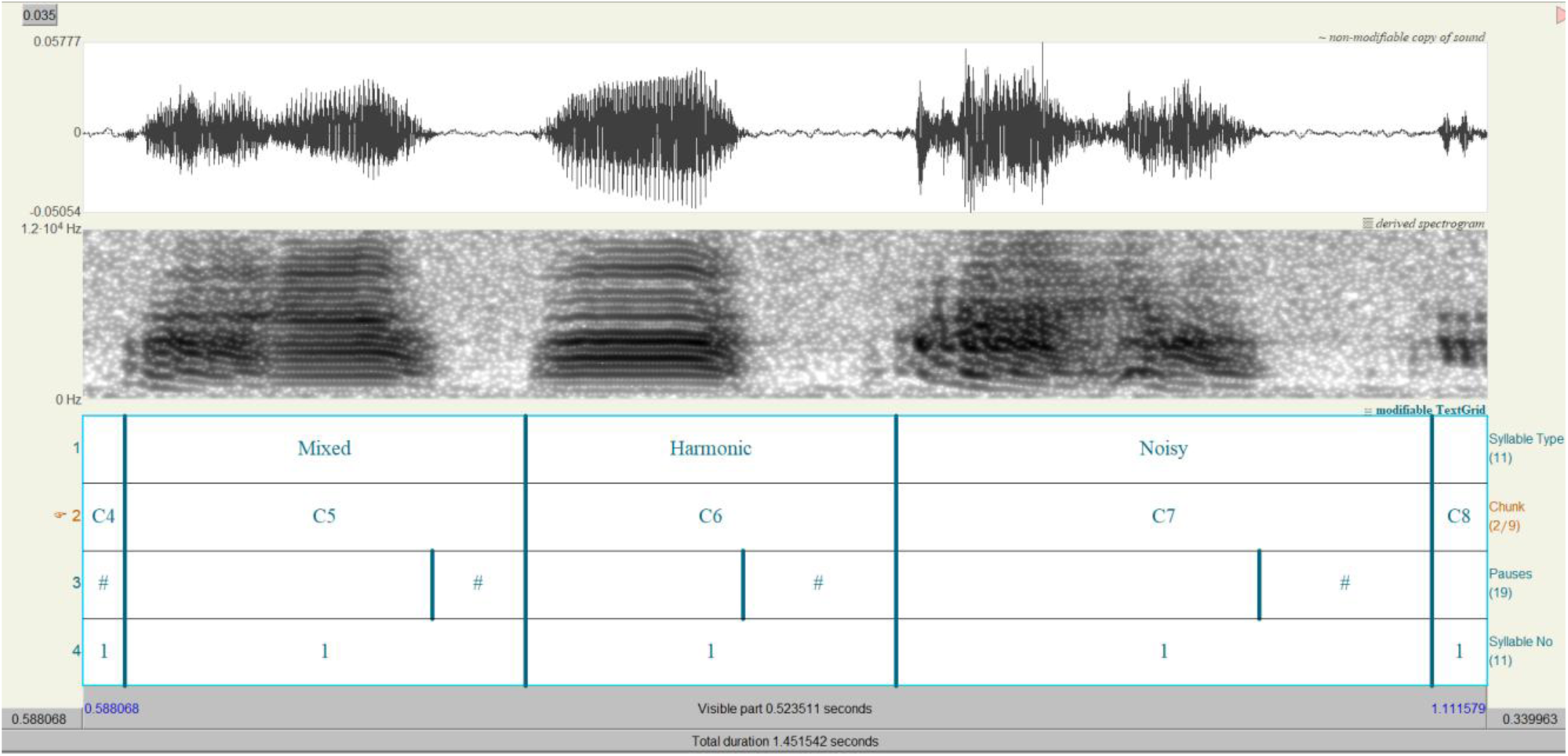
Example of harmonic, noisy, and mixed syllable segmentation in Praat. Example of syllables produced by the zebra finch Bk86 from older age group. From top to bottom: waveform, narrowband spectrogram (0-12 kHz), and annotation tiers. Annotation tiers are ordered as follows: (1) syllable type; (2) chunk number (C5-7) within the recording; (3) pauses (#); and (4) syllable count. In the narrowband spectrogram of the motif, it is possible to identify distinct acoustic patterns characteristic of the different song syllables produced by the zebra finch. The noisy syllable is characterized by the absence of well-defined harmonics and by the lack of organization into horizontal lines, reflecting a more chaotic sound without clear harmonic structure. Harmonic syllables, in contrast, exhibit relatively stable periodicity and are organized into horizontal bands. Mixed syllables combine features of both noisy and harmonic components. In these syllables, organized segments with harmonics aligned in horizontal bands coexist with disorganized segments in which the harmonics become less discernible.

In the aging dataset analyzed in Badwal *et al*. (2020), motifs and individual syllables were manually segmented in Praat version 6.4.07 (Boersma & Weenink, 2025). Syllables were subsequently classified into harmonic, noisy, and mixed categories (see Figure 1) based on spectrographic inspection (Badwal *et al*., 2019; Badwal *et al*., 2020), following established criteria (Badwal *et al*., 2019; Badwal *et al*., 2020; Gordon *et al*., 2025). Harmonic syllables were identified by the presence of clear, parallel harmonic stacks with stable periodicity and a well-defined *f*_o_; noisy syllables lacked harmonic organization and were dominated by broadband aperiodic energy; and mixed syllables contained both harmonic and noisy acoustic components within the same syllable. Classification was performed by one researcher and reviewed by a second experienced researcher (Badwal *et al*., 2020). Parameters related to *f*_o_ peaks (e.g., mean *f*_o_ peak bandwidth, *f*_o_ peak rate, *f*_o_ peak standard deviation, and *f*_o_ peak standard deviation timing) were not analyzed because only a limited number of reliable values could be extracted from the zebra finch vocalizations. These birds produce rapidly modulated songs with abrupt frequency transitions and frequent aperiodic segments, which reduce *f*_o_-tracking stability and compromise the automatic detection of well-defined *f*_o_ peaks. Consequently, the available data were considered insufficient for robust statistical analysis for these peak measurements.

In the aging dataset, the motifs of different UD singing birds were compared across age groups (younger, middle aged, and older). Birds were recorded during a single two-hour morning session and did not undergo any experimental manipulation.

In the original dataset from Medina *et al*. (2022), songs were segmented into individual motifs using the “Explore and Score” feature in SAP 2011. This procedure generated tables of syllable-level acoustic feature scores (e.g., *f*_o_, Wiener entropy, frequency modulation, and amplitude modulation). These scores were then used as input for the Vocal Inventory Clustering Engine (VoICE), a MATLAB-based tool that groups syllables into clusters based on multidimensional acoustic similarity, thereby identifying unique syllable types within each bird’s repertoire (Burkett *et al*., 2015). Syllables that were not reliably clustered into consistent modules were excluded from further analysis (Medina *et al*., 2022).

For the Medina *et al*. (2022) dataset, motif detection in the present study was performed using a custom MATLAB-based (MathWorks, 2026a) pipeline to automatically identify motifs within continuous recordings. First, a template motif was manually selected for each bird based on visual inspection of the spectrogram and waveform. Preference was given to renditions that exhibited the most consistent spectral structure and syllable timing, were free from overlapping noise, and occurred frequently throughout the session. Audio recordings were high-pass filtered at 200 Hz and normalized to reduce low-frequency noise and amplitude variability. The selected motif template was then compared against continuous recordings using normalized cross-correlation analysis. Motif occurrences were automatically identified based on correlation peaks exceeding a bird-specific similarity threshold (normalized cross-correlation coefficients ranging from 0.4 to 0.9 across individuals), with minimum peak-distance constraints applied to avoid duplicate detections of the same vocal sequence. Motif boundaries were subsequently established relative to the temporal position of the correlation peak and the duration of the original motif template.

Using this semi-automated pipeline, 25 motifs (see Figure 2) per bird were manually selected from the detected motifs for acoustic analysis, following our prior work that this provides sufficient power to detect experimental differences in juvenile and adult finches (Miller *et al*., 2010).

**Figure 2.**
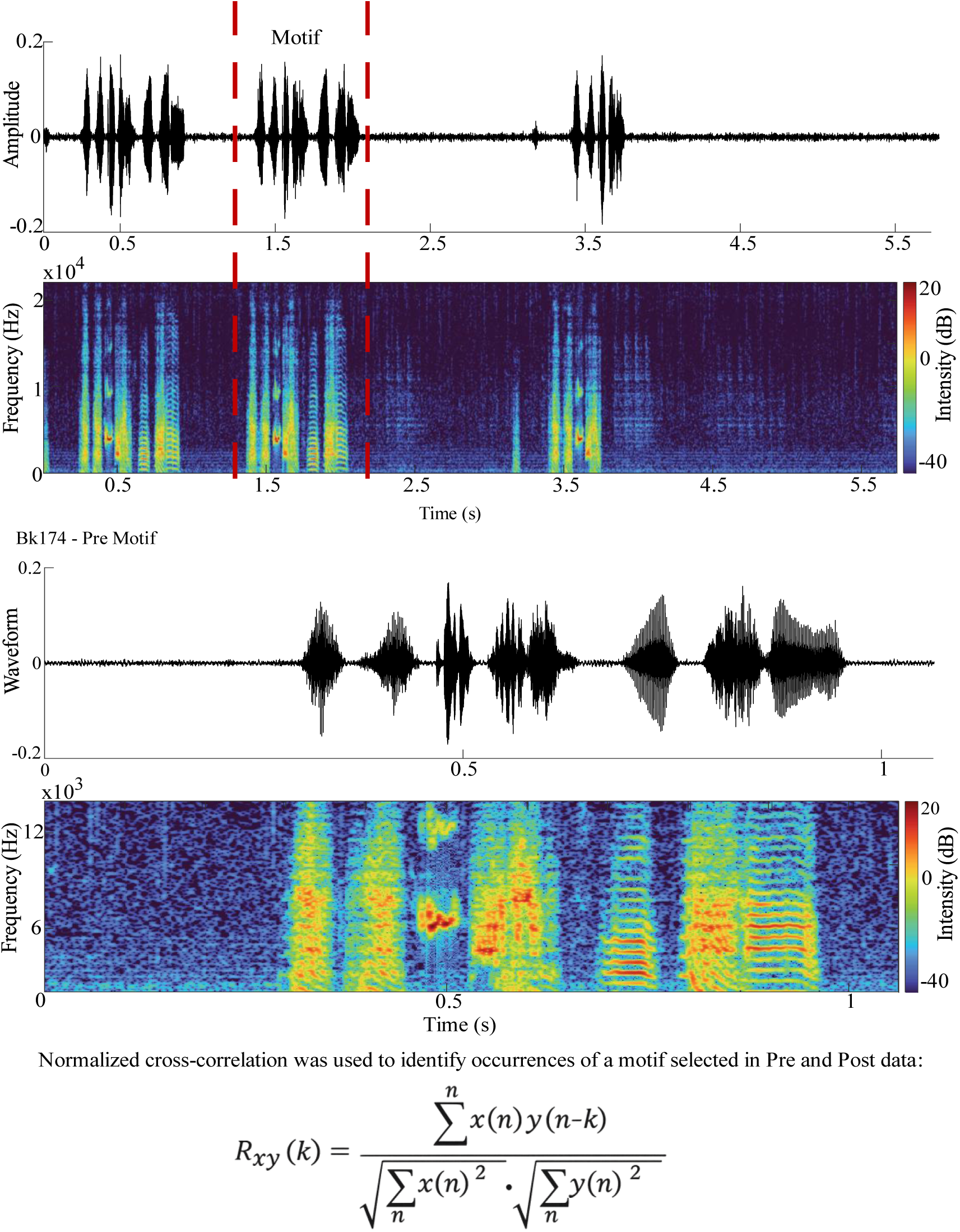
Motif extraction in MATLAB. Example of acoustic motif detection using normalized cross-correlation in bird Bk174 (bird identification number) prior to viral injection (Pre). The upper panel shows the waveform and spectrogram of a song segment containing multiple occurrences of the target motif. The first motif shown in the spectrogram was also identified as a valid match; however, the middle motif was selected for illustration purposes. The final motif was not selected because only a partial occurrence of the motif was present within the analyzed segment. The lower panel shows an expanded view of the selected motif (in the upper panel). The normalized cross-correlation function was used for acoustic motif detection, where R_xy_(k) denotes the normalized cross-correlation at lag k, x(n) represents the analyzed audio signal, y(n) represents the reference motif, and k corresponds to the temporal displacement between the two signals. Higher values of R_xy_(k) indicate greater similarity between the audio signal and the reference motif.

In the PD model dataset, the same birds were recorded before and after viral injection. For each bird, 25 motifs were analyzed pre-injection and 25 motifs were analyzed two months post-injection, selected across three recording mornings in both the ASYN and GFP groups, because previous work showed that vocal changes become evident two months after alpha-synuclein overexpression in Area X (Medina *et al*., 2022).

In the present study, for the Medina *et al*. (2022) dataset, introductory notes and calls were excluded from all analyses, and silent pauses were segmented by marking the offset of acoustic energy of one syllable and the onset of acoustic energy of the subsequent syllable.

### Acoustic analysis

Acoustic analysis was conducted using the Prosody Descriptor Extractor script in Praat (Barbosa, 2021), which extracts 25 parameters related to *f*_o_ (central and dispersion descriptors), as well as spectral, intensity, and temporal domains (see Table 1), using an *f*_o_ range of 400-2000 Hz for *f*_o_ detection. This range was adopted because the *f*_o_ of the zebra finch vocalizations is substantially higher than that of human voice: in humans, the spectral energy of harmonic syllables (e.g., vowels) is markedly attenuated above 4,000 Hz, whereas in zebra finches, attenuated harmonic energy can extend up to 10,000 Hz (Badwal *et al*., 2019). For the present analyses, parameter extraction was syllable-dependent. The *f*_o_-based parameters and voice-quality parameters were extracted exclusively from harmonic syllables, as only these syllables exhibit a continuous and reliable *f*_o_ contour in the spectrogram (see Figure 1).

**Table 1.** Systematic overview of the parameters.

| Parameter |  | Description | Unit |
| --- | --- | --- | --- |
| **Fundamental frequency ($f_0$)** | $f_{0med}$ | Median of $f_0$ | Hz/st re 1 Hz |
| | $f_{0sd}$ | Standard deviation of $f_0$ | Hz/st re 1 Hz |
| | $f_{0SAQ}$ | Semi-amplitude between $f_0$ quartiles | Hz/st re 1 Hz |
| | $f_{0min}$ | Minimum $f_0$ value | Hz/st re 1 Hz |
| | $f_{0max}$ | Maximum $f_0$ value | Hz/st re 1 Hz |
| | $f_{0peakwidth}$ | Mean bandwidth of $f_0$ peaks | Hz/st re 1 Hz |
| | $f_{0peak\_rate}$ | Rate of $f_0$ peaks | Peaks/second |
| | $sd f_{0peak}$ | Standard deviation of $f_0$ peaks | Peaks/second |
| | $df_{0posmean}$ | Mean of positive first derivative of $f_0$ | Hz/frame |
| | $df_{0negmean}$ | Mean of negative first derivative of $f_0$ | Hz/frame |
| | $sdt f_{0peak}$ | Standard deviation of $f_0$ peak positions over time | Seconds |
| | $df_{0sdpos}$ | Standard deviation of positive $f_0$ slope | Hz/frame |
| | $df_{0sdneg}$ | Standard deviation of negative $f_0$ slope | Hz/frame |
| **Related to Intensity** | emph | Spectral emphasis | dB |
|  | Mean Intensity | Mean Intensity | dB |
|  | cvint<sup>2</sup> | Coefficient of variation for intensity | % |
| **Voice Quality** | HNR | Harmonic-to-Noise Ratio | dB |
|  | sLTASmed | LTAS<sup>3</sup> slope at low frequencies | dB/frame |
|  | sLTAShigh | LTAS<sup>3</sup> slope at high frequencies | dB/frame |
|  | Jitter | Cycle-to-cycle variations in glottal periods | % |
|  | Shimmer | Cycle-to-cycle amplitude perturbation of glottal periods | % |
| **Rhythm** | Speech Rate | Number of syllabic units per second, including silent pauses | Syllables per second |
|  | Articulation Rate | Number of syllabic units per second, excluding silent pauses | Syllables per second |
| **Pause measures** | durSIL | Duration of silent pauses. | Milliseconds |
|  | IPI | Interpause Interval | Seconds |

<sup>1</sup>Frequency measures expressed in “st re 1 Hz” correspond to semitone values calculated relative to a reference frequency of 1 Hz using a logarithmic frequency scale; <sup>2</sup>Normalized measure, unaffected recording distance; <sup>3</sup>LTAS (Long-Term Average Spectrum) refers to the long-term spectral energy distribution of the acoustic signal across frequencies.

Intensity-related parameters were extracted for all syllables and analyzed separately by syllable type. In the original version of the Prosody Descriptor Extractor script (Barbosa, 2021), mean intensity was computed internally but remained uncorrected for chamber effects and was not written to the output table, as only the coefficient of variation for intensity (cvint) was exported. To measure average vocal intensity, we adapted the script to retain the computed mean intensity values (mint) and applied a chamber-effect correction factor prior to exporting the final data table. This correction was enabled by pre-calibration measurements using a 1,000-Hz reference tone, which compensated for acoustic artifacts associated with microphone position and individual chamber characteristics.

Temporal and pause-related parameters were computed independently of syllable type, as these measures do not rely on continuous *f*_o_ estimation. Given species-specific acoustic properties, spectral emphasis (emph) calculations were adapted from human speech conventions. In the original definition of emph, the low-band threshold is calculated as the mean *f*_o_ plus 1.5 standard deviations (Fant *et al*., 2000). In humans, this yields a threshold of approximately 400 Hz due to the lower *f*_o_ values of human voice. In zebra finches, however, the much higher *f*_o_ values of birdsong result in a substantially higher threshold, which more appropriately captures the distribution of energy across low- and high-frequency bands for zebra finch vocalizations.

## Statistical Analysis

All statistical analyses were conducted in RStudio version 2025.09. Because most acoustic parameters violated assumptions of normality and homoscedasticity, nonparametric statistical tests were used throughout, with the significance level set at α = 0.05. For all analyses, descriptive statistics (mean, median, and standard deviation) were computed from bird-level means for each condition and parameter.

For between-group comparisons in the aging dataset, effect size was estimated using epsilon-squared (ε^2^) derived from the Kruskal-Wallis test (Tomczak & Tomczak, 2014). For within-subject Pre-Post comparisons, effect magnitude was quantified using the standardized *r* statistic derived from the Wilcoxon Z value.

### Aging Dataset (Badwal *et al*., 2020): Between-Group Comparisons

As described above, between-group comparisons in the aging dataset were conducted using nonparametric statistical procedures. All analyses were performed separately for each acoustic parameter and, when applicable, for each motif chunk. However, for statistical testing, values from all motifs belonging to the same bird were first averaged, and these bird-level means were used as the independent observations.

For parameters related to *f*_o_, spectral, intensity, and temporal domains (excluding speech and articulation rates, analyzed separately), differences across age groups were assessed using the Kruskal-Wallis test. When a significant omnibus effect was detected, pairwise Wilcoxon rank-sum tests were performed with Holm correction for multiple comparisons to control Type-I error. Speech and articulation rates were analyzed separately due to their cumulative temporal nature. Bird-level values were used in Kruskal-Wallis and post hoc tests.

Statistical power analysis was conducted as a sensitivity-based post hoc estimation for all inferential tests. For Kruskal-Wallis analyses, statistical power was derived from ε^2^ effect sizes by converting ε^2^ to the corresponding Cohen’s *f*^2^ metric and estimating non-centrality parameters under the F-distribution framework (Cohen, 1988). This procedure provided an estimate of the probability of detecting group differences given the observed effect sizes, sample sizes, and number of groups. Power estimates were computed for each acoustic-prosodic parameter at the bird-level aggregation, and should be interpreted as descriptive indicators of test sensitivity rather than as inferential evidence of effect presence or absence.

### Pre-Post Comparisons (Medina *et al*., 2022): Within-Subject Design

For longitudinal comparisons between pre- and two-month post-injection conditions, within-subject analyses were conducted using the Wilcoxon signed-rank test on bird-level mean values, due to non-normality and repeated-measures structure.

Effect sizes were quantified using the standardized *r* statistic derived from the Wilcoxon Z value (*r* = Z divided by the square root of N), also interpreted according to Cohen (1988) benchmarks for the correlation coefficient *r*: 0.10-0.30 small, 0.30-0.50 medium, and ≥0.50 large.

Speech and articulation rates were analyzed at the bird level by summing values across motifs within each session prior to paired tests. Statistical power was estimated from observed *r* values and sample sizes, with detailed procedures provided in the Supplemental Material.

## Results

Results are presented separately for the aging and PD datasets. We first describe age-related differences across the adult lifespan and subsequently examine pre-post changes following alpha-synuclein overexpression (ASYN) or GFP control injections in song nucleus Area X. Detailed descriptive statistics, inferential tests, and effect sizes are provided in Tables S1-S3.

### Aging dataset: Non-linear age trajectories in spectral and temporal acoustic features

Many *f*_o_, spectral, and intensity parameters show non-linear age-related trajectories: *f*_o_ measures drop at middle age, while spectral measures (slLTASmed, slLTAShigh, and emph) and the coefficient of variation for intensity (cvint) increase in older birds, and HNR dips at middle age before recovering.

In the aging dataset, significant age-related differences were observed across several acoustic domains. For harmonic syllables, which showed the most numerous effects, *f*_o_ measures (*f*_o_med, *f*_o_min, and *f*_o_max) dropped at middle aged compared to younger adults, with no increase in older birds (see Figure 3). Pairwise comparisons (see Table S2) confirmed that middle aged birds differed significantly from both younger and older groups for these parameters, whereas younger and older birds did not differ. Spectral measures (slLTASmed and slLTAShigh) and the cvint increased markedly in older birds for the harmonic syllables, with effect sizes (ε^2^) of 0.137, 0.135, and 0.209, respectively, for slLTASmed, slLTAShigh and cvint (see Figures 4-5). These effect sizes were substantially larger than those observed for *f*_o_ measures: *f*_o_med (ε^2^ = 0.024), *f*_o_max (ε^2^ = 0.024), *f*_o_min (ε^2^ = 0.024), *f*_o_sd (ε^2^ = 0.012), *f*_o_SAQ (ε^2^ = 0.009), and d*f*_o_negmean (ε^2^ = 0.008). Mean intensity was also higher in middle aged and older birds compared to younger (see Figure 5). HNR dipped at middle age and returned to high values in older birds, while emph was highest in older birds (see Table S1).

**Figure 3.**
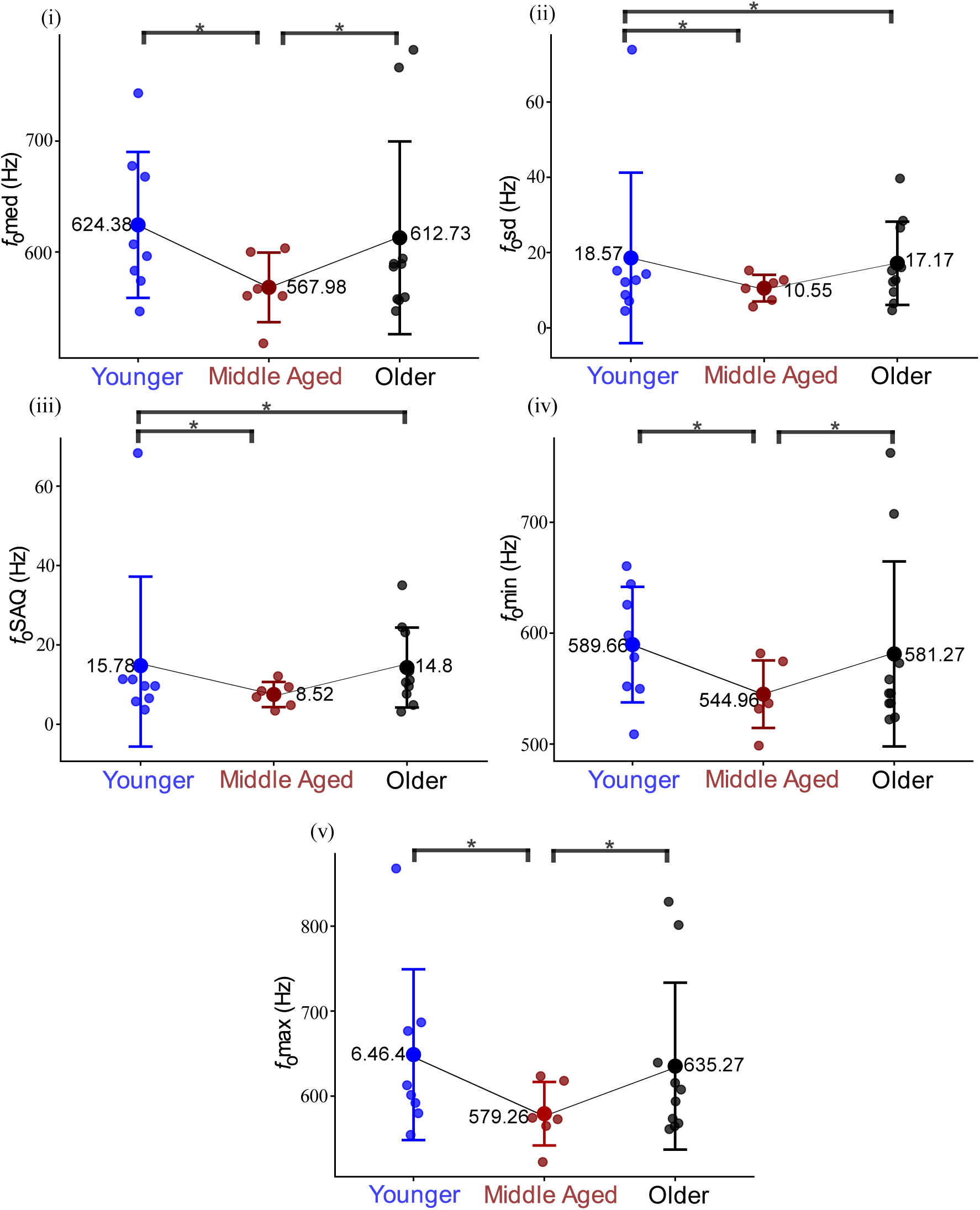
*f*_o_-related parameters for aging groups. Comparison of fundamental frequency measures across younger (n = 8), middle aged (n = 8), and older (n = 10) zebra finches. Graphics (i-v) show (i) median of *f*_o_ (*f*_o_med), (ii) standard deviation of *f*_o_ (*f*_o_sd), (iii) semi-amplitude between *f*_o_ quartiles (*f*_o_SAQ), (iv) minimum of *f*_o_ (*f*_o_min), and (v) maximum of *f*_o_ (*f*_o_max). The x-axis indicates age group (Younger, Middle aged, and Older birds), and the y-axis shows the corresponding parameter values in Hertz (Hz). Individual dots represent values from a single bird, large circles indicate group means, and error bars represent ±1 standard deviation (SD). Statistical significance was assessed using pairwise Wilcoxon rank-sum tests with Holm correction following significant Kruskal-Wallis effects. * Indicates significant between-group differences (p < 0.05).

**Figure 4.**
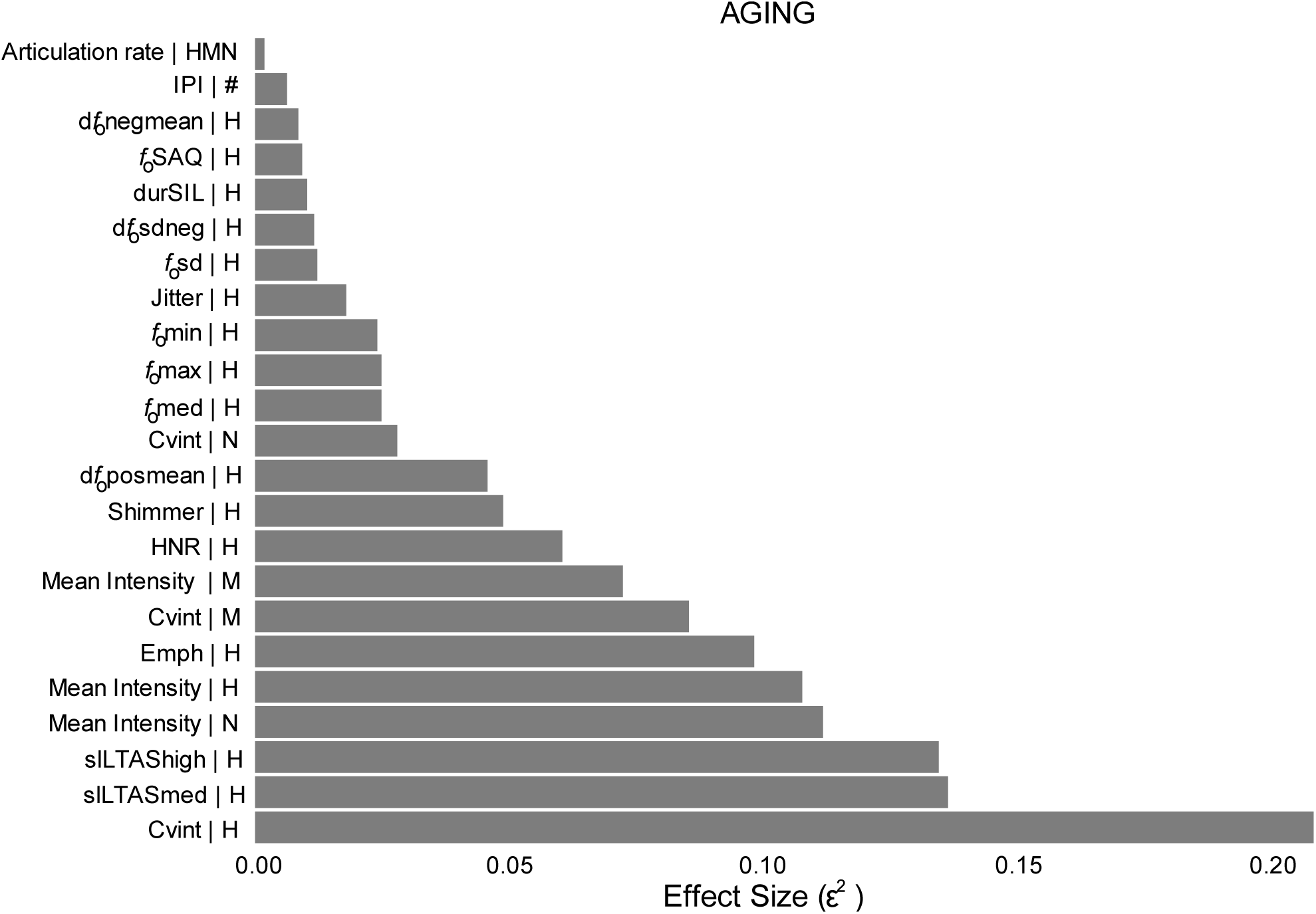
Effect size for aging groups according to syllable type

**Figure 5.**
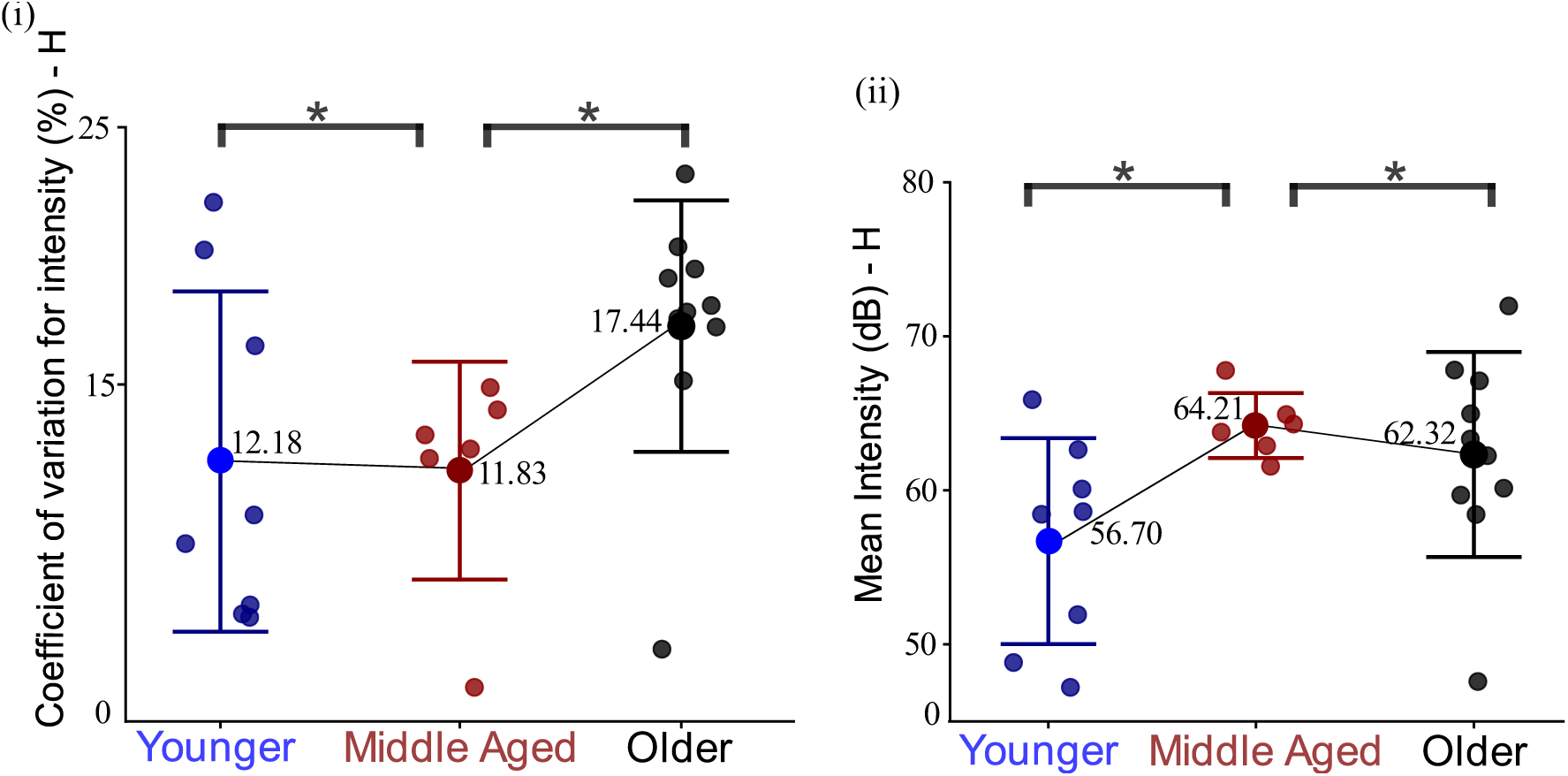
Coefficient of variation for intensity (cvint) and Mean intensity (mint) for harmonic syllables (H) across age groups. Individual and group-level differences in acoustic-prosodic measures are shown for younger, middle aged (mage), and older birds. Small colored points represent individual bird means (each point corresponds to the average of all syllable tokens per bird within the harmonic syllable category). Large open circles indicate group means for each age condition, and error bars represent ±1 standard deviation (SD) across birds. * Indicate statistically significant differences between-groups (p < 0.05), based on pairwise comparisons.

For mixed and noisy syllables, significant effects were fewer and smaller, primarily in intensity-related parameters (see Table 2 and Figure 4). No significant age differences were found for speech rate or articulation rate (see Table 2).

**Table 2.**
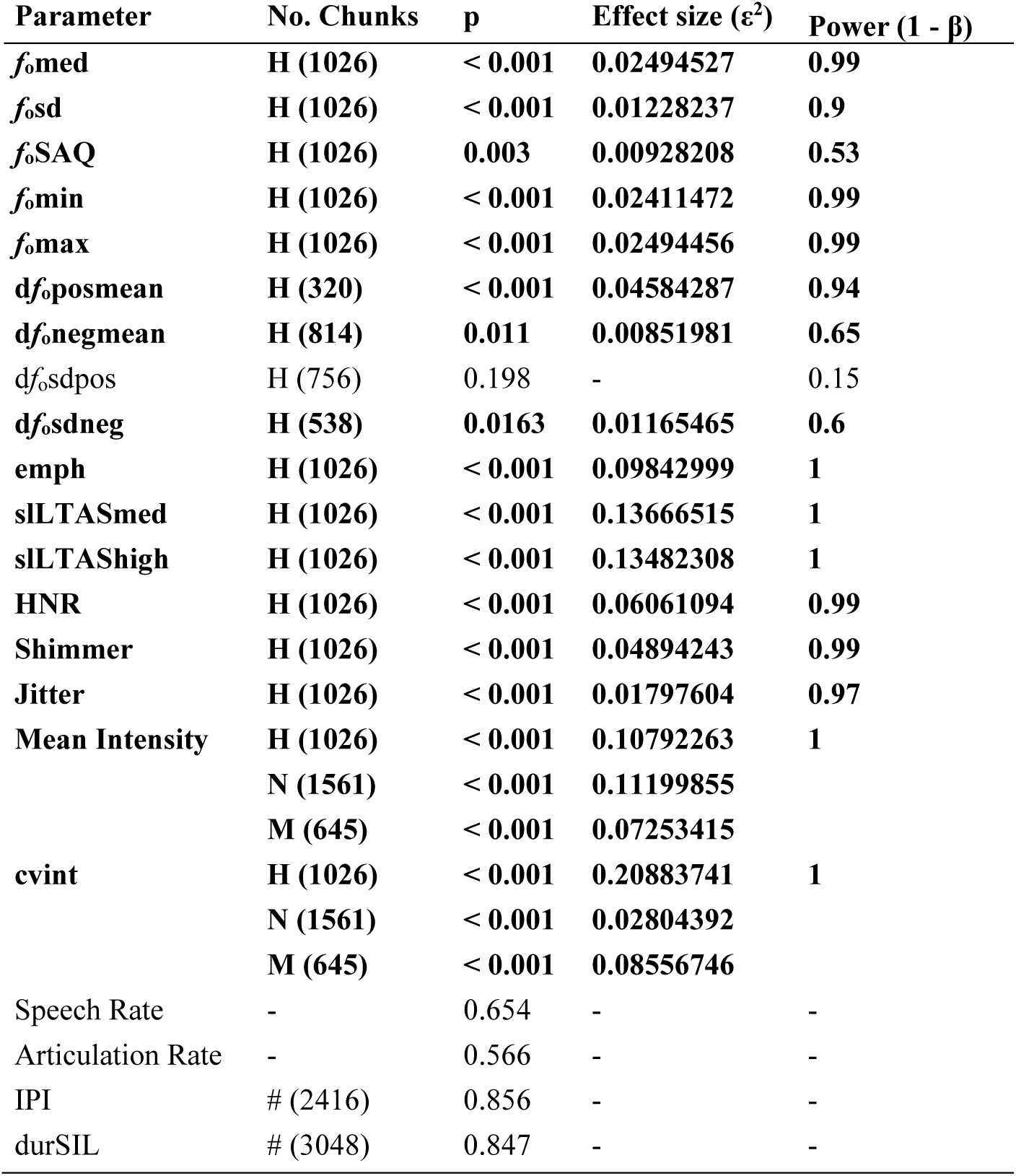
Effect sizes for age-related differences in acoustic-prosodic parameters. Values correspond to Kruskal-Wallis tests comparing younger, middle aged, and older birds. Effect size is reported as ε^2^, which estimates the proportion of variance in ranked data attributable to group differences. Analyses were conducted per motif chunk and, when applicable, separately for harmonic (H), noisy (N), and mixed (M) syllables. Statistical power (1 - β) was estimated from the primary Kruskal-Wallis analysis for each parameter and is reported once per parameter. For measures analyzed across multiple syllable types (e.g., Mean Intensity and cvint), power estimates are shown only for the primary analysis and were not recalculated separately for each syllable subtype. Post hoc comparisons were performed using Wilcoxon rank-sum tests with Holm correction. Parameters with statistically significant differences (α = 0.05) are shown in bold. Speech and Articulation Rates measures are calculated at the motif level.

Effect sizes revealed that the largest age-related effects were concentrated in spectral measures and intensity variability, indicating that these acoustic features were more sensitive to aging than fundamental frequency measures. These findings expand upon Badwal *et al*. (2020) which identified age-related changes in CPPS, intensity, and song rate for all syllable types, by examining a broader set of acoustic-prosodic measures and reporting effect sizes that were not previously analyzed.

Statistical power was generally high for the parameters showing the largest age-related effects, particularly cvint, slLTASmed, slLTAShigh, mean intensity, and emph (1 - β ≥ 0.99). In contrast, several fundamental frequency measures exhibited smaller effect sizes and more variable power estimates, indicating that age-related changes were most robustly expressed in spectral features and intensity-related measures.

Among harmonic syllables, significant age-related effects were observed across multiple acoustic domains. *f*_o_ measures (*f*_o_med, *f*_o_sd, *f*_o_SAQ, *f*_o_min, and *f*_o_max) showed significant effects, although generally of small magnitude. Larger effects were observed for intensity- and spectral-related measures, including cvint (ε^2^ = 0.209), slLTASmed (ε^2^ = 0.137), slLTAShigh (ε^2^ = 0.135), mean intensity (ε^2^ = 0.108), and emph (ε^2^ = 0.098).

Significant effects were also identified for voice-quality measures, including HNR, shimmer, and jitter, although with smaller effect sizes.

For mixed syllables, significant age-related effects were primarily observed for intensity-related measures, including mean intensity (ε^2^ = 0.073) and cvint (ε^2^ = 0.086).

For noisy syllables, significant age-related effects were likewise concentrated in intensity-related measures, including mean intensity (ε^2^ = 0.112) and cvint (ε^2^ = 0.028).

### Larger pre-post acoustic effect sizes following alpha-synuclein overexpression

In the PD dataset, significant pre-post differences were observed in both the ASYN and GFP control groups (see Table 3 and Table S3). Many significant findings were shared across groups, but some parameters differed. Significant changes in emph (harmonic syllables) were observed in both groups, although the effect was larger in the ASYN group. Significant changes in mean intensity (mixed syllables) were observed only in the ASYN group, whereas significant changes in *f*_o_med (harmonic syllables) and cvint (mixed syllables) were observed only in the GFP group. Speech rate and articulation rate did not differ significantly between recording sessions in either group. Given the many overlapping findings between the two groups, we sought to determine next if the two groups could be differentiated by the magnitude of the changes.

**Table 3.** Effect sizes for group differences in acoustic-prosodic parameters in PD dataset. Values correspond to effect sizes (*r*) derived from paired Wilcoxon signed-rank tests comparing pre- and post-injection conditions. Analyses were conducted per motif chunk. ASYN = alpha-synuclein overexpression group; GFP = control group expressing green fluorescent protein. H = harmonic syllables; N = noisy syllables; M = mixed syllables. Pause-related measures (IPI and durSIL) are indicated by (#) and were computed independently of syllable type. Post hoc comparisons were performed using the Wilcoxon rank-sum test with Holm correction. Parameters with statistically significant differences (α = 0.05) are shown in bold. Speech and Articulation Rates measures are calculated at the motif level.

| Parameter | Group | Syllable | p | Size effect ($r$) | Power ($1 - \beta$) |
| --- | --- | --- | --- | --- | --- |
| $f_0$med | ASYN | H | 0.218 | 0.022697 | 0.83 |
|  | GFP | H | < 0.001 | **0.143238** | **1** |
| $f_0$sd | ASYN | H | < 0.001 | **0.255149** | **1** |
|  | GFP | H | < 0.001 | **0.127701** | **1** |
| $f_0$SAQ | ASYN | H | < 0.001 | **0.262756** | **1** |
|  | GFP | H | < 0.001 | **0.112836** | **1** |
| $f_0$min | ASYN | H | **0.002** | **0.011823** | **0.33** |
|  | GFP | H | < 0.001 | **0.160667** | **1** |
| $f_0$max | ASYN | H | < 0.001 | **0.041172** | **0.99** |
|  | GFP | H | < 0.001 | **0.106990** | **1** |
| df<sub>0</sub>posmean | ASYN | H | 0.167 | 0.030088 | 0.3 |
|  | GFP | H | 0.583 | 0.013801 | 0.07 |
| df<sub>0</sub>negmean | ASYN | H | < 0.001 | **0.261146** | **1** |
|  | GFP | H | < 0.001 | **0.311499** | **1** |
| df<sub>0</sub>sdpos | ASYN | H | < 0.001 | **0.813389** | **1** |
|  | GFP | H | < 0.001 | **0.787914** | **1** |
| df<sub>0</sub>sdneg | ASYN | H | < 0.001 | **0.396393** | **1** |
|  | GFP | H | < 0.001 | **0.119622** | **0.99** |
| slLTASmed | ASYN | H | < 0.001 | **0.132774** | **1** |
|  | GFP | H | < 0.001 | **0.107365** | **1** |
| slLTAShigh | ASYN | H | < 0.001 | **0.110554** | **1** |
|  | GFP | H | < 0.001 | **0.114236** | **1** |
| HNR | ASYN | H | < 0.001 | **0.075540** | **1** |
|  | GFP | H | < 0.001 | **0.179981** | **1** |
| Shimmer | ASYN | H | < 0.001 | **0.067309** | **1** |
|  | GFP | H | < 0.001 | **0.150156** | **1** |
| Jitter | ASYN | H | **0.004** | **0.107747** | **1** |
|  | GFP | H | < 0.001 | **0.159507** | **1** |
| emph | ASYN | H | < 0.001 | **0.045703** | **0.99** |
|  | GFP | H | **0.041** | **0.042205** | **0.96** |
| Mean Intensity | ASYN | H | < 0.001 | **0.302497** | **1** |
|  | ASYN | M | < 0.001 | **0.321028** | **1** |
|  | ASYN | N | < 0.001 | **0.0918** | **0.99** |
|  | GFP | H | < 0.001 | **0.054369** | **0.99** |
|  | GFP | M | **0.226** | **0.154** | **1** |
|  | GFP | N | < 0.001 | **0.123** | **1** |
| cvint | ASYN | H | < 0.001 | **0.119159** | **1** |
|  | ASYN | M | **0.003** | **0.276** | **1** |
|  | ASYN | N | < 0.001 | **0.102** | **1** |
|  | **GFP** | **H** | **0.037** | **0.176344** | **1** |
|  | **GFP** | **M** | **< 0.001** | **0.332** | **1** |
|  | **GFP** | **N** | **< 0.001** | **0.093** | **1** |
| Speech Rate | ASYN | - | 0.286 | 0.375154 | 0.51 |
|  | GFP | - | 1 | 0.042796 | 0.025 |
| Articulation rate | ASYN | - | 1 | 0.019744 | 0.025 |
|  | GFP | - | 0.833 | 0.128388 | 0.053 |
| IPI | ASYN | # | **< 0.001** | **0.18733438** | **1** |
|  | **GFP** | **#** | **< 0.001** | **0.1311177** | **1** |
| durSIL | ASYN | # | **< 0.001** | **0.022559** | **1** |
|  | **GFP** | **#** | **< 0.001** | **0.071179** | **1** |

Figures 6 and 7 (see also Table 3) summarize effect sizes for pre-post comparisons across acoustic domains and syllable types in the ASYN and GFP groups, respectively. Both groups showed significant differences across many parameters, with standard deviation of positive *f*_o_ slope (d*f*_o_sdpos) exhibiting the largest effect size in both groups. However, effect size analysis revealed clearer differences between the ASYN and GFP groups than comparisons based on statistical significance alone. The ASYN group showed medium effect sizes for standard deviation of *f*_o_ (*f*_o_sd), Semi-amplitude between *f*_o_ quartiles (*f*_o_SAQ_)_, and standard deviation of negative *f*_o_ slope (d*f*_o_sdneg) whose values increased following alpha-synuclein overexpression (see Figure 6). In the GFP control group, these parameters had small effect sizes and mean values decreased. The ASYN group also showed medium effect sizes for mean intensity and cvint in the harmonic and mixed syllable types, whereas the GFP control group showed small effect sizes for these measures.

**Figure 6.**
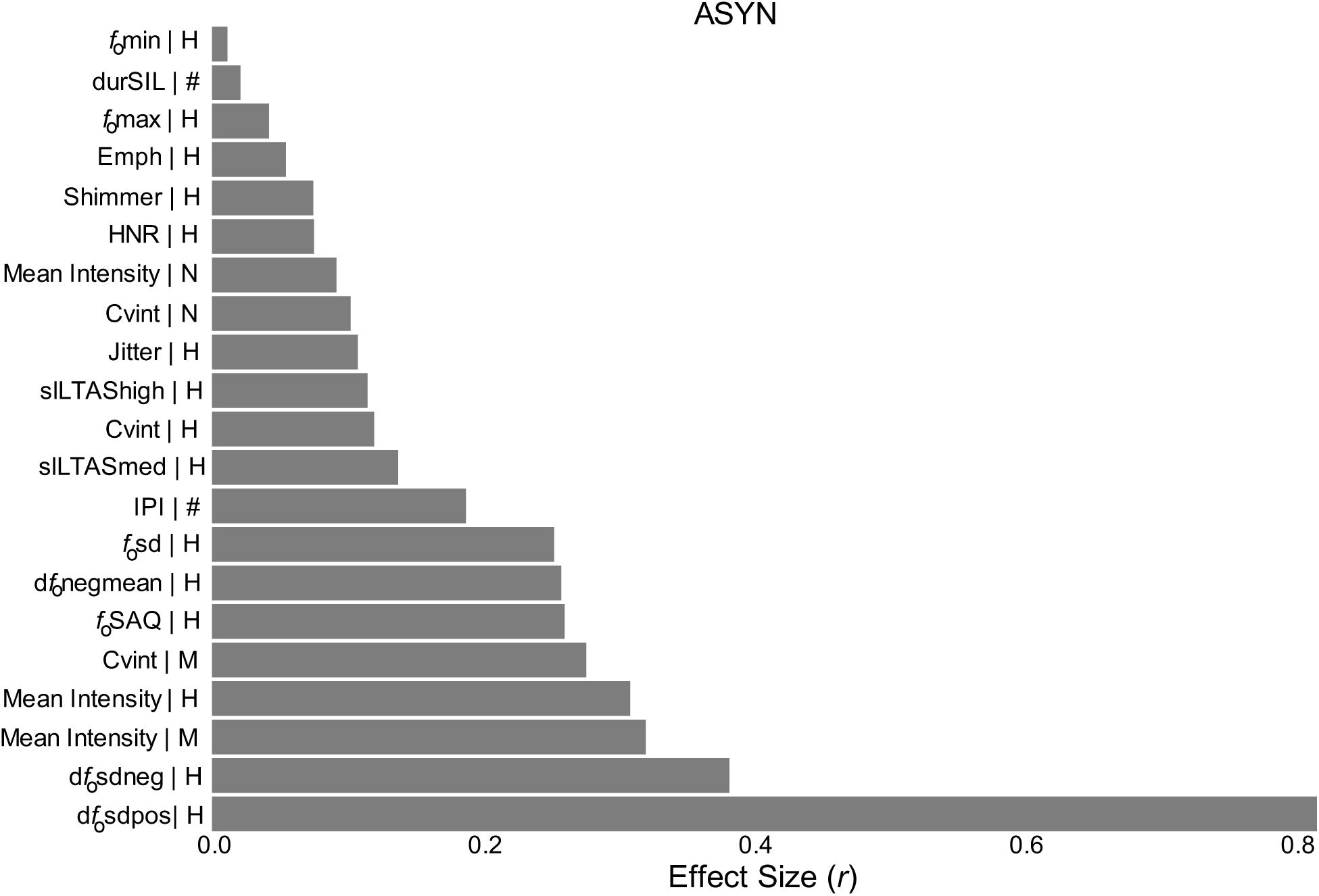
Effect size for ASYN group (Pre vs. Post) according to syllables type

**Figure 7.**
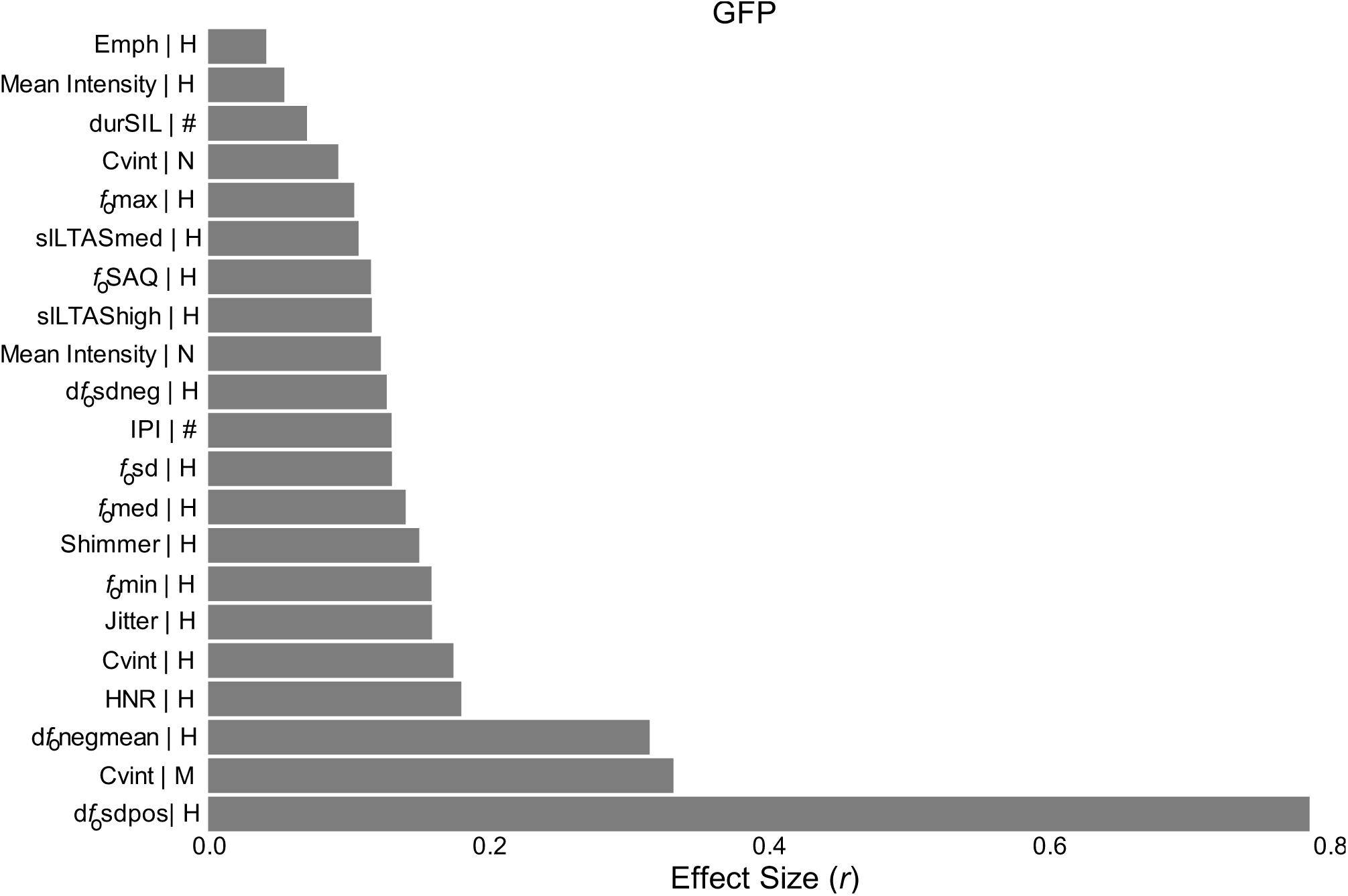
Effect size for GFP group (Pre vs. Post) according to syllables type

The GFP group generally showed smaller effects, with the largest changes observed for d*f*_o_sdpos and cvint (see Figures 6-7). Representative examples of *f*_o_ variability and intensity-related measures for both groups are presented in Figure S1.

Power analyses indicated high sensitivity for most acoustic parameters showing significant pre-post effects, particularly those related to *f*_o_ variability, intensity, spectral slope, and pause structure (1 - β ≥ 0.99, Table 3). In contrast, the non-significant findings for speech rate and articulation rate were accompanied by relatively low power estimates, indicating that the absence of detectable effects for these temporal measures should be interpreted cautiously.

## Discussion

### Non-linear trajectories of vocal aging evident in birdsong

The present study investigated how normative aging and alpha-synuclein overexpression in a PD model influence zebra finch vocal production using a broad set of acoustic-prosodic measures commonly employed in human voice research. Overall, both aging and alpha-synuclein overexpression were associated with alterations across overlapping acoustic domains, with intensity-related measures showing consistent sensitivity to vocal variation across datasets. However, age-related differences encompassed multiple acoustic domains, including *f*_o_, spectral, voice quality, and intensity measures, whereas alpha-synuclein overexpression was associated with a more selective pattern involving vocal intensity and specific aspects of *f*_o_ regulation, particularly d*f*_o_sdneg.

### *f*_o_ measures drop in middle age and more stable patterns emerge in older finches

Although significant age-related differences were observed across multiple *f*_o_ measures, the largest effect sizes (ε^2^) were observed for spectral and intensity-related parameters (emph, slLTASmed, slLTAShigh, mean intensity, and cvint). These measures were also associated with the highest statistical power estimates, suggesting that these parameters provided the most sensitive measures for detecting age-related differences in the present sample.

These findings should be interpreted in light of the parameter-specific analytical framework adopted. *f*_o_, spectral, and voice quality measures were extracted exclusively from harmonic syllables because reliable periodicity is required for these analyses, whereas intensity measures were evaluated across all syllable types. Within this framework, harmonic syllables showed age-related changes across multiple acoustic domains, while mixed and noisy syllables primarily exhibited alterations in vocal intensity. Although zebra finch syllables cannot be directly equated with human speech sounds, harmonic syllables present acoustic properties analogous to vowel-like segments, particularly stable periodic structure that permits *f*_o_ estimation, whereas noisier syllables contain acoustic features more comparable to consonant-like elements.

This distribution of effects suggests that age-related variation is expressed across multiple acoustic properties of harmonic syllables, whereas changes in mixed and noisy syllables were largely restricted to vocal intensity. Harmonic syllables may offer greater sensitivity for detecting subtle age-related changes in vocal motor control, while intensity appears to represent a more general marker across syllable types.

Voice quality measures (HNR, shimmer, and jitter) also showed significant age-related effects, though generally smaller in magnitude (see Figure 4). In contrast, global temporal measures (speech rate, articulation rate, and pause metrics) remained unchanged across age groups, with no statistically significant differences. This dissociation indicates that aging-related vocal changes are not uniformly distributed across acoustic domains. Notably, age-related changes did not follow a single directional pattern; *f*_o_ measures showed a striking non-linear, U-shaped pattern (see Figure 3), with middle aged birds showing lower *f*_o_ values and differing significantly from both younger and older groups, whereas younger and older birds showed comparable levels. This pattern suggests that *f*_o_ aging may reflect a transitional phase of altered vocal control during middle age followed by relative stabilization in older birds (Badwal *et al*., 2020; Higgins *et al*., 2025), while spectral and intensity-related measures showed more pronounced changes in older birds.

Importantly, non-linear trajectories of vocal aging, including U-shaped patterns of *f*_o_-related parameters, have also been reported in humans (Stathopoulos *et al*., 2011; Valente *et al*., 2024). These findings suggest that age-related changes in vocal frequency may not follow a uniformly progressive trajectory across species. The present findings therefore provide further evidence that vocal aging in zebra finches may involve transient changes during middle adulthood followed by relative stabilization in later life, although the specific mechanisms underlying these trajectories may differ across species.

Consistent with these findings, human voice aging research demonstrates that intensity and spectral energy distribution measures may be more sensitive to age-related changes than timing or speech rate measures (Schultz *et al*., 2023). Measures related to spectral energy distribution, voice turbulence, cepstral intensity, and frequency variability are among the strongest predictors of chronological age (Schultz *et al*., 2023).

From a physiological standpoint, spectral and intensity changes in aging voices reflect interactions among the respiratory system, laryngeal biomechanics, and central motor control processes. Normal aging involves structural and functional changes to the respiratory apparatus and laryngeal tissues (decreases in lung elasticity, weakening of respiratory musculature, and changes in vocal fold viscoelastic properties) which affect airflow and phonatory coordination (Rojas *et al*., 2020; Cavallaro *et al*., 2024). These mechanisms provide plausible physiological frameworks for interpreting the prominence of spectral slope, intensity, and variability effects observed in zebra finch aging, although the specific contributions of respiratory, peripheral, and neural factors require direct investigation.

### Preservation of temporal parameters with aging

One of the most consistent findings was the relative preservation of global temporal organization despite significant changes in several acoustic domains. Pause-related measures (IPI and durSIL) were not significant, whereas speech and articulation rates showed only subtle age-related differences, with a negligible effect size for articulation rate. This pattern suggests that aging affects acoustic execution more strongly than the temporal organization of learned vocal sequences (see Table 2 and Table S2).

The relative preservation of global timing aligns with human aging research, where changes in speech rate and timing are not consistently associated with normative aging alone. Temporal characteristics such as speech rate, articulation rate, and pause frequency vary more with speech style than with age itself in cross-sectional comparisons of younger and older adults, suggesting that relatively stable temporal organization may reflect the influence of compensatory or contextual factors (Taylor *et al*., 2020).

In songbirds, temporal aspects of song structure appear relatively stable across much of adulthood. Although song timing undergoes maturation during early life stages in zebra finches (Glaze & Troyer, 2012), evidence for large age-related changes in adult timing is limited. Our earlier cross-sectional study revealed that mean song and articulation rates at the motif level were faster in middle aged birds than in younger and older adults (Badwal *et al*., 2020). Evidence from other species is mixed: Bengalese finches show increased inter-syllable durations in older birds (Cooper *et al*., 2012), while others report increased stereotypy with age (James & Sakata, 2014).

Together with the present findings, this body of work suggests that normative aging primarily affects fine-grained acoustic features related to vocal execution rather than the temporal organization of learned vocal sequences. The stereotyped and highly sequential nature of birdsong may constrain large-scale temporal changes, may contribute to maintaining stability in timing even as spectral, intensity, and other fine-grained acoustic features exhibit age-related variability. Understanding how aging affects both brain networks (Higgins *et al*., 2025) and the vocal organ is an important direction for future research in the zebra finch model. Our findings strengthen its value as an experimentally tractable model for investigating the biological basis of vocal aging.

### Effect sizes reveal larger changes in *f*_o_ variability, vocal intensity, and frequency modulation following alpha-synuclein overexpression compared to controls

A notable finding was that substantial pre-post acoustic changes occurred in both ASYN and GFP birds, suggesting that longitudinal factors (two-month recording, viral transduction, surgical intervention, or age-related changes in young adults) could have impacted vocal behavior independently of alpha-synuclein overexpression.

Despite these shared effects, some acoustic measures exhibited larger changes in ASYN than in GFP. Among harmonic syllables, descending frequency modulation variability (d*f*_o_sdneg) showed a markedly larger effect in ASYN (*r* = 0.396) than in GFP (*r* = 0.120). Mean intensity also showed substantially larger effect sizes in ASYN than in GFP. Measures of *f*_o_ variability (*f*_o_sd and *f*_o_SAQ) exhibited approximately two-fold larger effect sizes in ASYN (*r* = 0.255-0.263) than in GFP (*r* = 0.113–0.128), with mean values rising in ASYN versus falling in controls, suggesting greater alterations in vocal control in the ASYN condition.

The clearest differences between ASYN and GFP were in harmonic syllables, where mean intensity and d*f*_o_sdneg exhibited substantially larger effect sizes following alpha-synuclein overexpression. In contrast, ascending frequency modulation variability (d*f*_o_sdpos) showed similarly large effect sizes in both groups (ASYN: *r* = 0.81; GFP: *r* = 0.79), suggesting stronger influence from longitudinal or procedural factors than from alpha-synuclein-related mechanisms. Overall, effect size patterns indicate that vocal intensity and descending frequency modulation were most strongly associated with alpha-synuclein overexpression, whereas *f*_o_ variability measures exhibited effect sizes approximately two-fold greater in ASYN than GFP.

Several parameters (spectral measures, voice quality measures, and pause-related parameters) showed significant changes in both groups with comparable effect sizes, suggesting non-specific effects. Larger effects for mean intensity, d*f*_o_sdneg, and the two-fold increases in *f*_o_sd and *f*_o_SAQ may be more specifically associated with alpha-synuclein overexpression.

Such fine-grained changes are consistent with the role of basal ganglia circuitry in regulating movement initiation, motor scaling, and behavioral variability. In songbirds, neural activity within cortico-basal ganglia circuits contributes directly to acoustic variability and adaptive vocal modification (Woolley & Kao, 2015). Experimental manipulations of these circuits can alter *f*_o_ regulation, vocal variability, and song adaptation (Saravanan *et al*., 2019). Similarly, in humans, basal ganglia dopaminergic dysfunction underlies PD and contributes to hypokinetic dysarthria, characterized by reduced vocal loudness, restricted *f*_o_ modulation, diminished prosodic variability, imprecise articulation, and variable speech timing (Moya-Galé & Levy, 2019).

Although the effects in ASYN birds were selective rather than widespread, measures associated with *f*_o_ variability and vocal intensity showed larger effect sizes in the alpha-synuclein group, suggesting that alpha-synuclein overexpression may preferentially influence fine-scale vocal motor regulation rather than global song structure.

### Intensity regulation as a shared vulnerability in aging and PD

Intensity-related measures ranked among the strongest effects in both datasets, regardless of whether the source was normative aging or longitudinal viral manipulation. In the aging dataset, mean intensity and cvint were among the most strongly affected parameters. In the PD dataset, significant pre-post differences were observed for both measures in ASYN and GFP, with generally larger effects in ASYN. These findings indicate that intensity-related measures are particularly sensitive to changes in vocal motor function across different biological contexts.

In human PD, reduced vocal intensity and increased variability of vocal effort are hallmarks of hypokinetic dysarthria, attributed to impaired integration between motor commands and sensory feedback, resulting in deficient motor scaling and reduced calibration of vocal output (Cavallaro *et al*., 2024). Speakers with PD often underestimate their vocal loudness and exhibit diminished capacity to generate and maintain adequate sound pressure levels, reflecting dysfunction within basal ganglia-cortical loops involved in gain control and motor amplitude regulation.

The present findings suggest that intensity control is a particularly sensitive marker of disruption in the vocal motor system, spanning both normative aging and disease-related conditions. Underlying mechanisms are likely multifactorial. In normative aging, alterations in vocal intensity and its variability have been linked to changes in respiratory support, pulmonary pressure generation, and vocal tissue biomechanics (Ramig *et al*., 2008). In the PD dataset, intensity-related changes in both ASYN and GFP indicate that longitudinal and procedural factors also contribute. Nevertheless, larger effects in ASYN are consistent with alpha-synuclein overexpression further disrupting neural mechanisms involved in motor scaling and vocal output control.

This partial overlap highlights intensity regulation as a shared vulnerability of the vocal system. Rather than providing a disease-specific marker, intensity-related measures may reflect the combined influence of peripheral biomechanical factors, respiratory function, and central neural control mechanisms. The consistent sensitivity of these parameters across both datasets suggests that vocal intensity may represent a useful indicator of vocal system integrity, while differences in effect magnitude may help distinguish normative aging from pathology-related changes.

### Implications for comparative and translational vocal research

Our findings reveal that many parameters are affected by normative aging, but a more restricted set showed larger changes following alpha-synuclein overexpression compared with controls, contributing to a growing body of work supporting the use of songbirds as models for human vocal motor disorders. The aging dataset showed that spectral and intensity-related measures were among the strongest correlates of vocal aging, whereas the PD dataset indicated that alpha-synuclein-related effects were more selectively concentrated in descending frequency modulation, *f*_o_ variability, and larger changes in vocal intensity relative to controls. The partial overlap between aging- and alpha-synuclein-related acoustic alterations underscores the importance of accounting for normative aging when evaluating vocal biomarkers in both humans and animal models. Measures of *f*_o_ variability and intensity regulation emerged as candidate acoustic features for monitoring vocal motor alterations associated with aging and alpha-synuclein -related dysfunction.

Future work combining acoustic analyses with electrophysiological recordings from song-control nuclei, respiratory musculature, and syringeal muscles (Adam *et al*., 2021) will help to disentangle the neural and peripheral mechanisms underlying vocal changes associated with both aging and alpha-synuclein overexpression.

Beyond characterizing age- and disease-related vocal changes, this study demonstrates that acoustic-prosodic measures from human voice research can be applied to zebra finch song to capture biologically meaningful variation linked to normative aging and alpha-synuclein overexpression. The findings underscore the need to disentangle aging-related, longitudinal, and disease-related sources of vocal variability when evaluating candidate biomarkers. Collectively, these findings support the zebra finch as a valuable model for investigating vocal aging and neurodegenerative dysfunction, and highlight the potential of multidimensional acoustic analyses for identifying cross-species signatures of vocal motor change.

### Limitations and future directions

Some limitations should be acknowledged. Analyses were restricted to undirected song. Because female-directed and undirected songs differ in social, motivational, and neural substrates, future studies should examine whether aging- and alpha-synuclein-related vocal changes generalize across behavioral contexts.

Although zebra finches provide a valuable model for studying learned vocal behavior, important limitations should be considered when drawing parallels with human aging. Aging in zebra finches occurs over a substantially shorter lifespan and under different developmental, physiological, and ecological conditions than in humans. In addition, birdsong does not capture the linguistic and semantic complexity of human speech. Direct comparisons should therefore be interpreted cautiously. Rather than serving as a direct model of human speech aging, zebra finches are best viewed as a comparative system for investigating general principles of vocal motor control, neural plasticity, and age-related changes in learned vocal behavior.

Despite these constraints, the present study offers a comprehensive and biologically grounded characterization of vocal changes associated with aging and alpha-synuclein overexpression in zebra finches. By integrating analytical approaches from human phonetics and animal bioacoustics, this work advances a comparative framework for investigating how vocal motor control is shaped by normative aging and neurodegenerative processes.

## Acknowledgements

The authors acknowledge the researchers responsible for the publicly available datasets analyzed in this study and their commitment to data sharing and open science practices, which made the present work possible.

## Data Availability Statement

All recordings were obtained from two open-access datasets collected from adult male zebra finches in the Miller colony at the University of Arizona between 2015 and 2020. The original datasets are freely accessible here: University of Arizona ReData repository:

- Miller, J. E., Samlan, R. A., Borgstrom, M. C., & Badwal, A. (2020). Raw birdsong data for “Middle Age, a Key Time Point for Changes in Birdsong and Human Voice.”. University of Arizona Research Data Repository. Media. https://doi.org/10.25422/azu.data.12020763;
- Medina, C. A., Vargas, E., Munger, S., & Miller, J. E. (2022). Raw birdsong data for “Vocal changes in a zebra finch model of Parkinson’s Disease characterized by alpha-synuclein overexpression in the song-dedicated anterior forebrain pathway”. University of Arizona Research Data Repository. Dataset. https://doi.org/10.25422/azu.data.16619782.v1

Processed data, analysis scripts, and supplementary materials associated with the present study are available through the University of Arizona ReData repository upon publication: Dal’Ava, Lucas Manca; Barbosa, Plinio; Miller, Julie Elizabeth (2026). Acoustic-Prosodic Biomarkers of Aging and Parkinson’s Disease in a Songbird. University of Arizona Research Data Repository. Dataset. https://doi.org/10.25422/azu.data.32015082

## Author contributions

Dal’Ava, Lucas M., University of Campinas (UNICAMP) and State University of São Paulo “Júlio de Mesquita Filho” (UNESP): Conceptualization, Data Curation, Formal Analysis, Investigation, Methodology, Software, Validation, Visualization, Writing-Original Draft, Writing-Review & Editing. Barbosa, Plinio A., University of Campinas (UNICAMP): Conceptualization, Methodology, Supervision, Validation, Writing-Review & Editing. Miller, Julie E., University of Arizona: Conceptualization, Methodology, Resources, Supervision, Validation, Writing-Review & Editing.

## Supplemental Material

Supplementary descriptive statistics and pairwise comparisons are provided in Tables S1, S2, and S3. Table S1 presents descriptive statistics of acoustic-prosodic measures by age group in the aging dataset. Table S2 summarizes pairwise post hoc comparisons between age groups in the aging dataset. Table S3 summarizes pre- and post-injection acoustic-prosodic measures for the ASYN and GFP groups in the PD dataset.

**Table S1.** Descriptive statistics of acoustic-prosodic measures by age group. Values are mean (SD). No. Chunks = number of analyzed motif chunks. H = harmonic; N = noisy; M = mixed syllables. *f*_o_-related and voice quality parameters extracted from harmonic syllables only; intensity parameters analyzed separately by syllable type; temporal and pause measures computed independently of syllable type. Cv = coefficient of variation.

**Table S2.** Pairwise post hoc comparisons between age groups in the aging dataset. P-values adjusted using Holm correction. Significant comparisons (α = 0.05) in bold.

**Table S3.** Descriptive acoustic-prosodic measures pre- and post-injection for ASYN and GFP groups. Values are mean and standard deviation. ASYN = alpha-synuclein overexpression; GFP = control. H, N, M = syllable types. Pause measures (IPI and durSIL) indicated by (#).

**Figure S1.** Representative *f*_o_ variability and intensity measures for harmonic syllables. Pre- to post-injection changes in ASYN and GFP groups. Graphics show (i) *f*_o_sd, (ii) *f*_o_SAQ, (iii) d*f*_o_sdpos, (iv) d*f*_o_sdneg, (v) mean intensity (mint) for harmonic syllables, and (vi) coefficient of variation for intensity (cvint) for harmonic syllables. Points = individual bird means; large points = group means; error bars = ±1 SEM. * p < 0.05.

## Supplemental Methods

Detailed procedures for statistical power estimation in pre-post comparisons.

## Supplemental Material

**Table S1.**
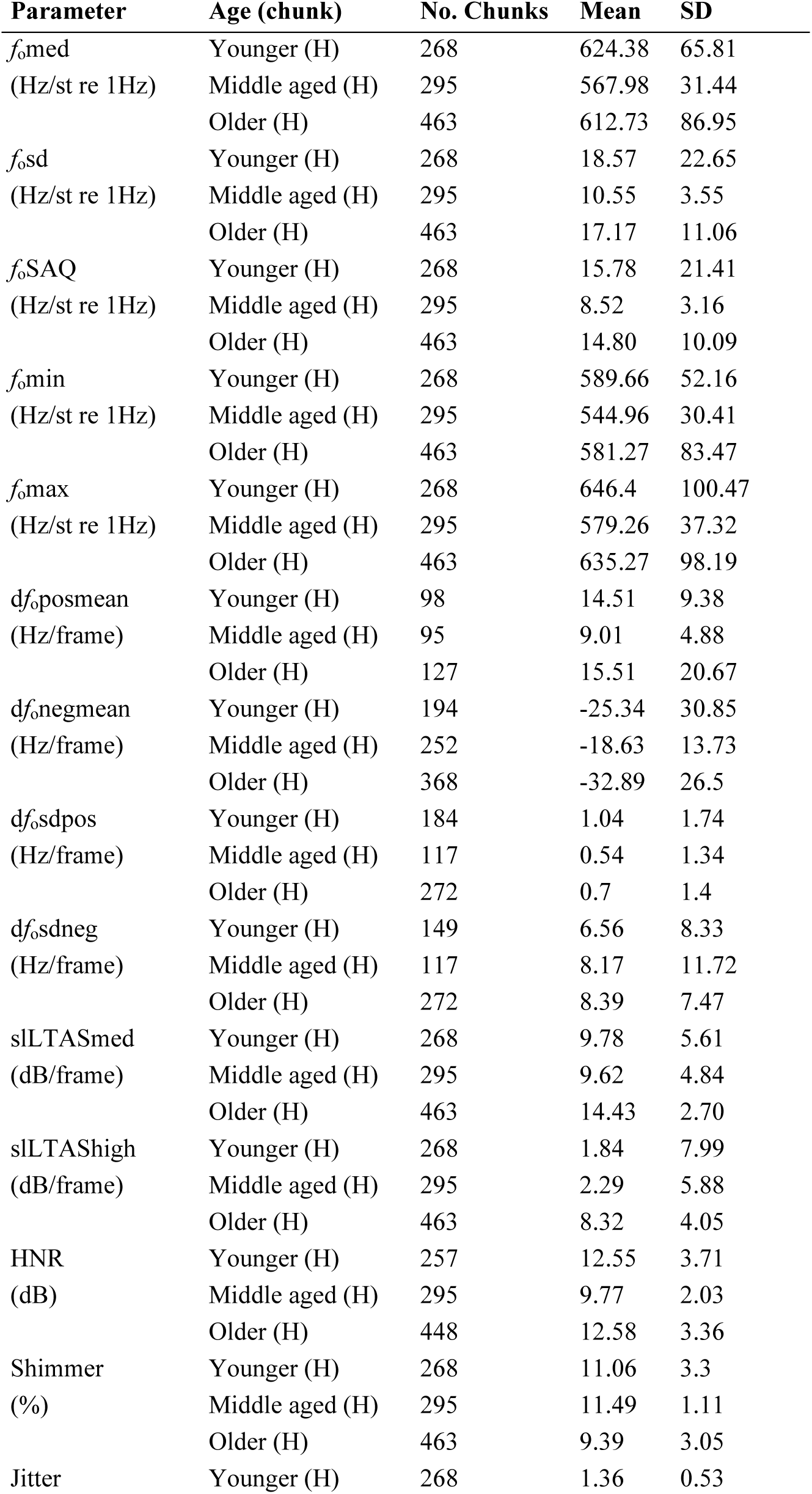
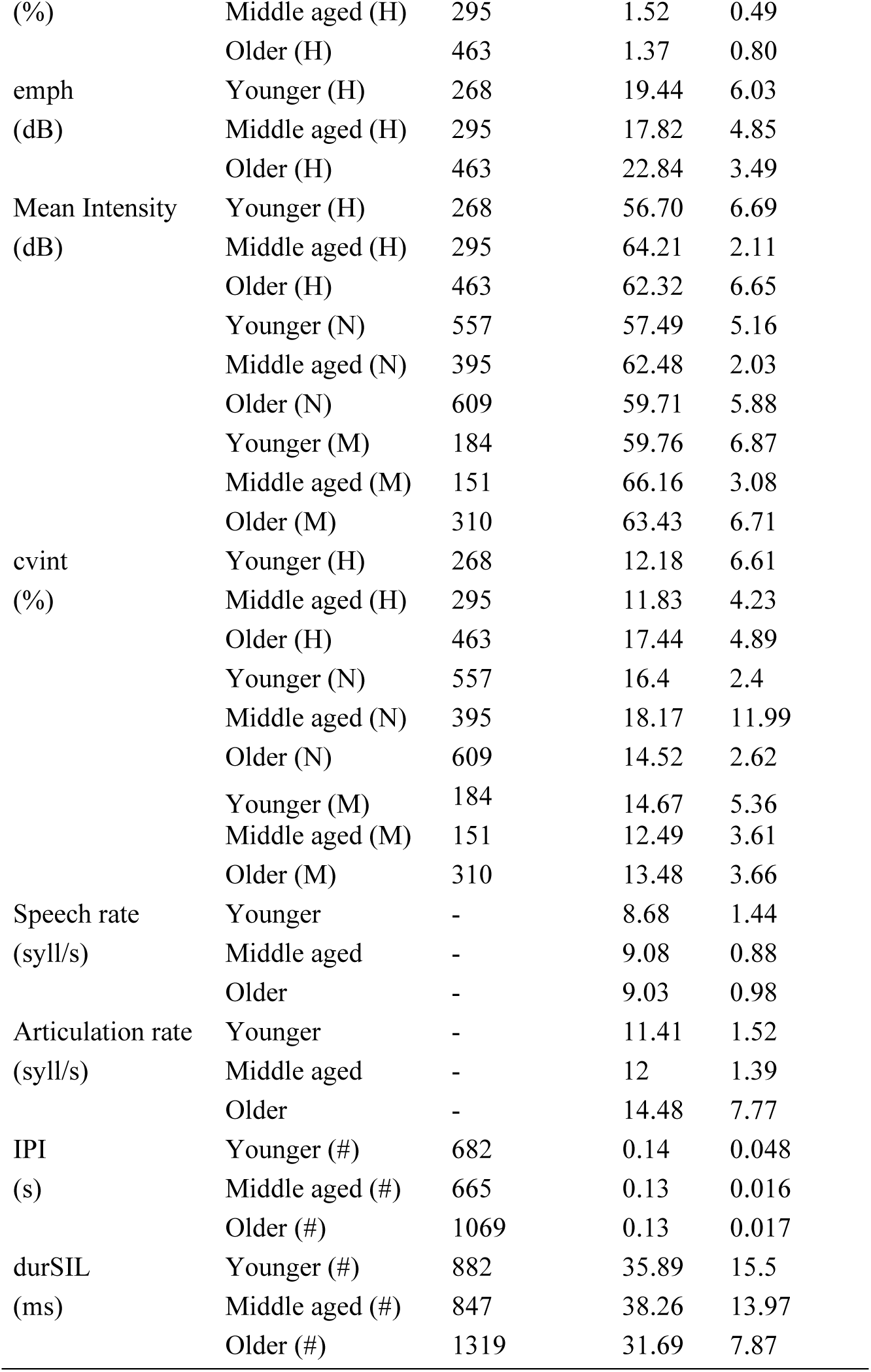
Descriptive statistics of acoustic-prosodic measures by age group. Values are reported as mean and standard deviation (SD). No. Chunks refers to the number of analyzed motif chunks contributing to each estimate. H = harmonic syllables; N = noisy syllables; M = mixed syllables. *f*_o_-related and voice quality parameters were extracted exclusively from harmonic syllables. Intensity (int) parameters were analyzed separately by syllable type, whereas temporal and pause measures were computed independently of syllable classification. Cv = coefficient of variation. Speech and Articulation Rates measures are calculated at the motif level.

**Table S2.**
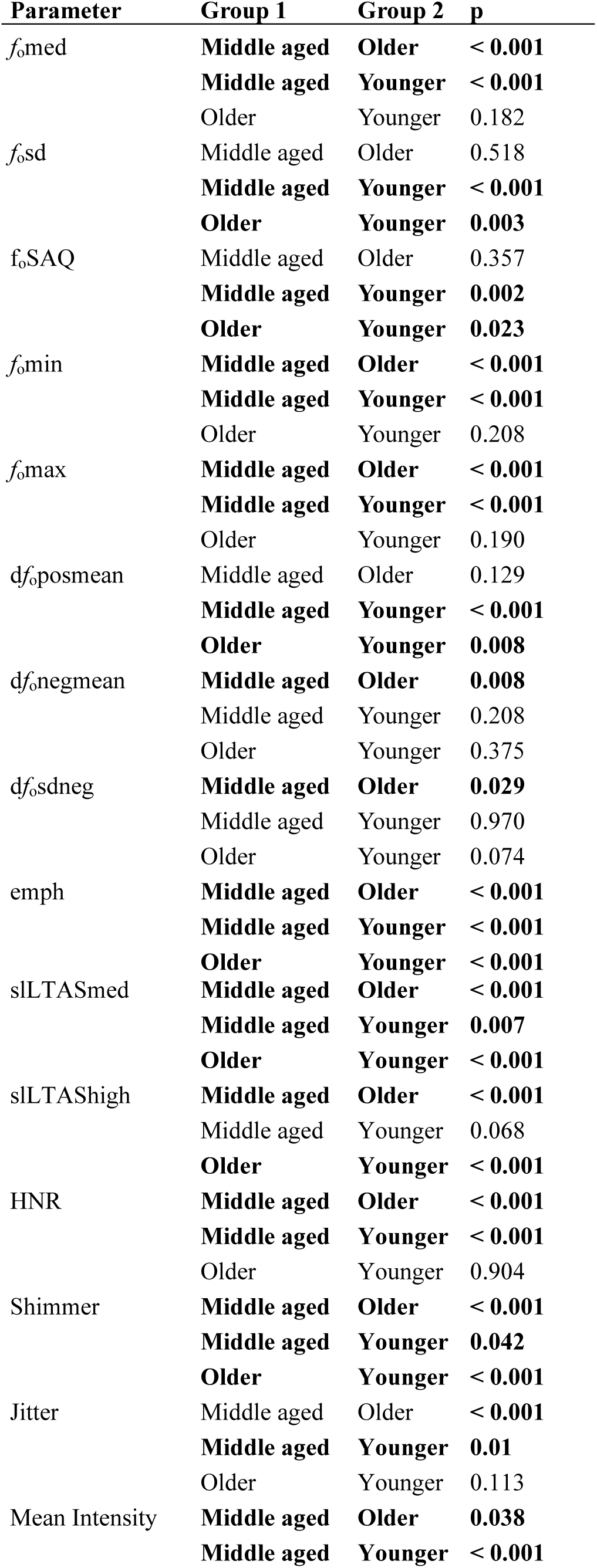
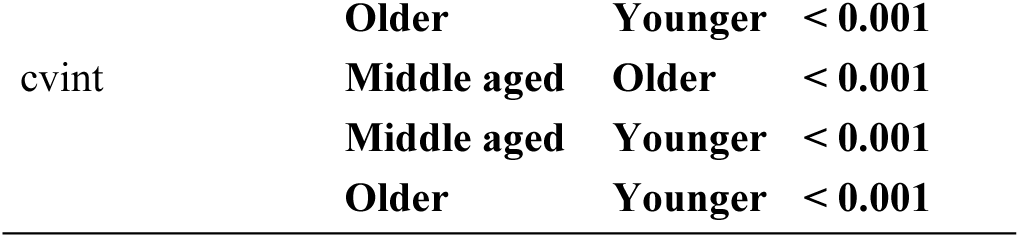
Pairwise post hoc comparisons between age groups in the aging dataset. Pairwise mean comparisons were performed using Wilcoxon rank-sum tests following significant Kruskal-Wallis effects. P-values were adjusted using the Holm correction for multiple comparisons. Only comparisons reaching statistical significance (α = 0.05) are shown in bold.

**Table S3.**
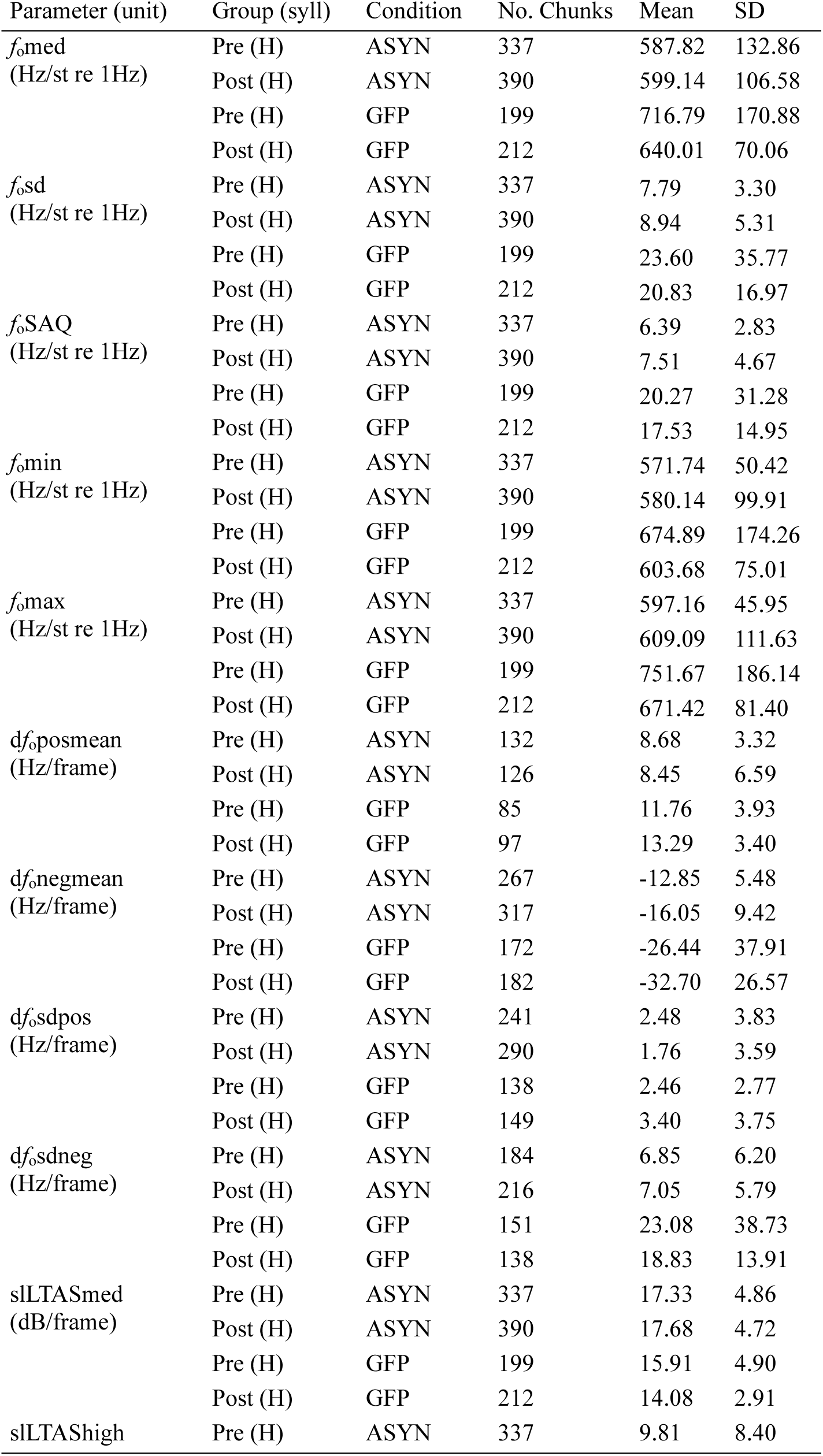
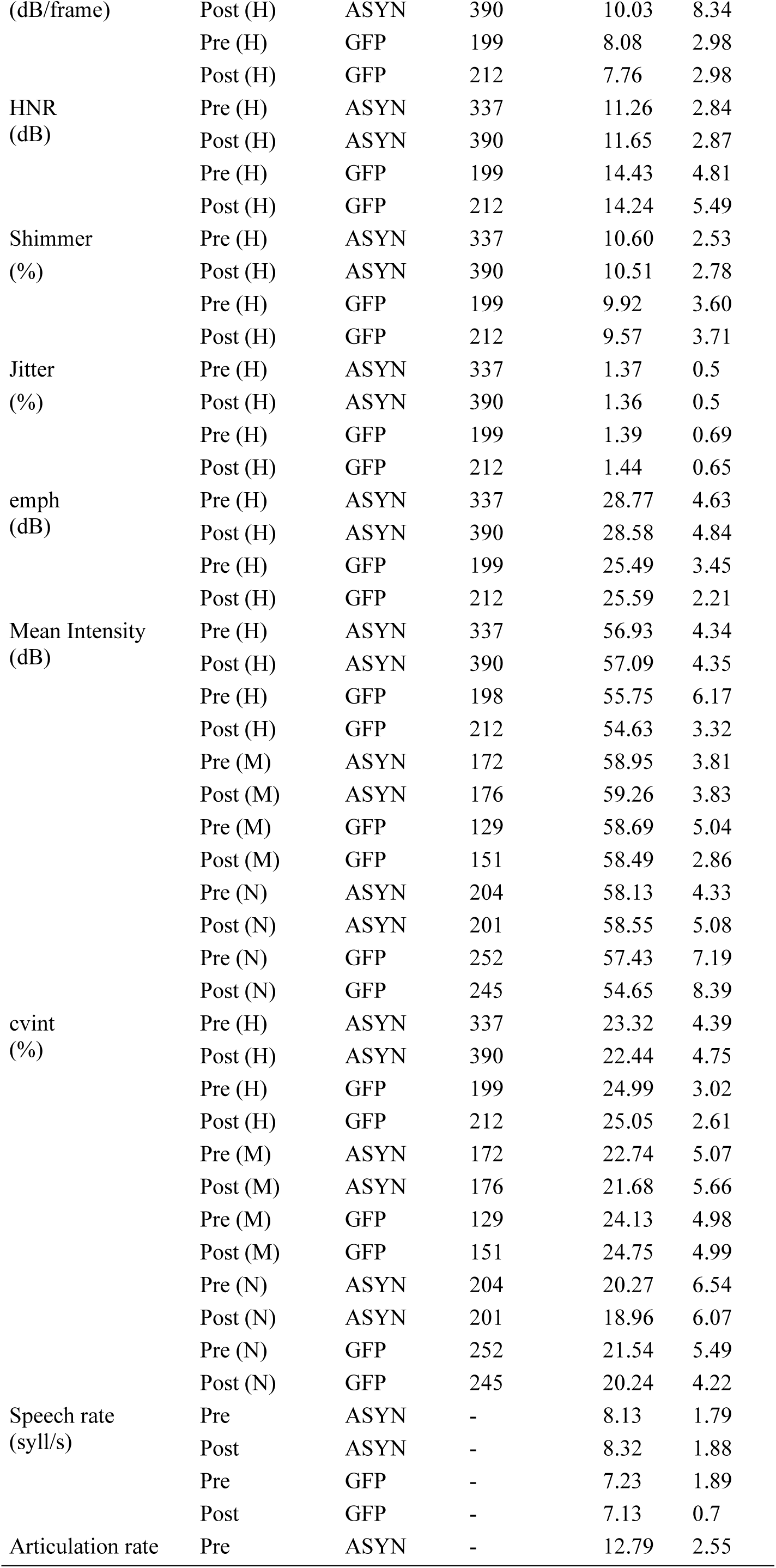
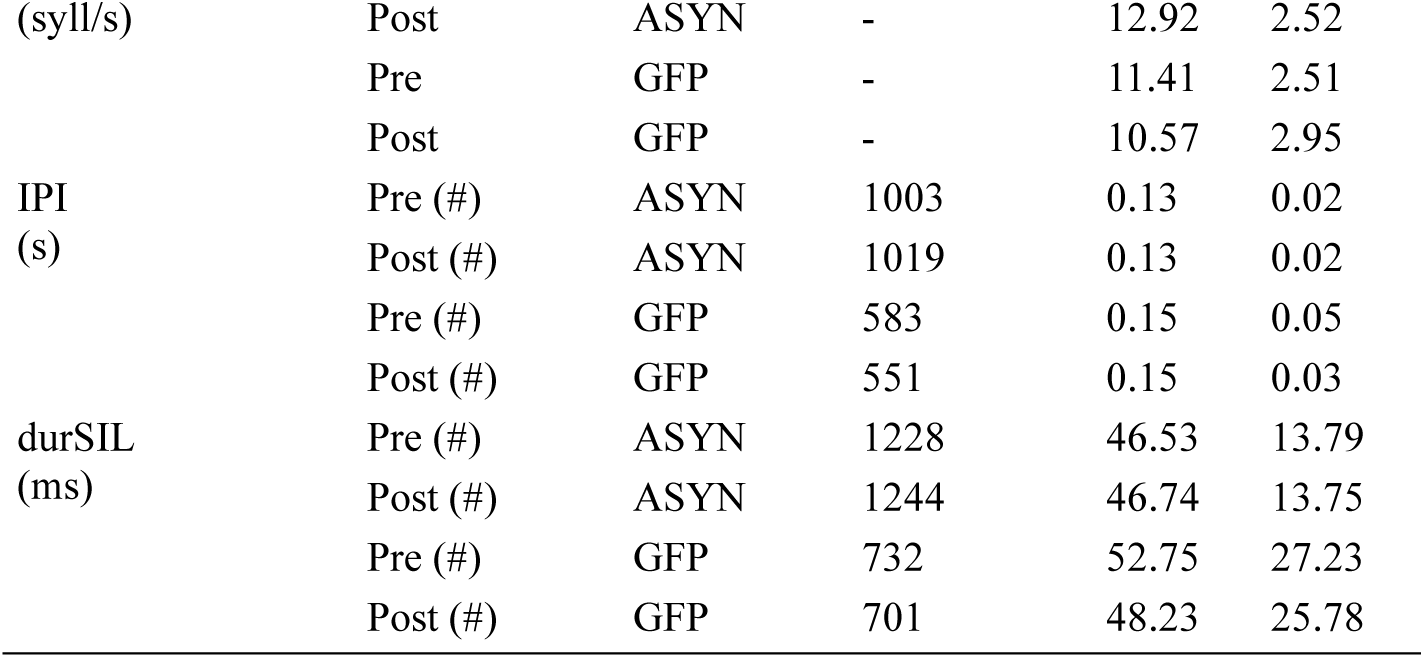
Descriptive acoustic-prosodic measures pre- and post-injection ASYN/GFP. Values are reported as mean and standard deviation (SD). No. Chunks indicates the number of analyzed motif chunks contributing to each condition. Pre and Post refer to recordings obtained before and after viral injection, respectively. ASYN = alpha-synuclein overexpression group; GFP = control group expressing green fluorescent protein. H = harmonic syllables; N = noisy syllables; M = mixed syllables. Pause-related measures (IPI and durSIL) are indicated by (#) and were computed independently of syllable type. Speech and Articulation Rates measures are calculated at the motif level.

**Figure S1.**
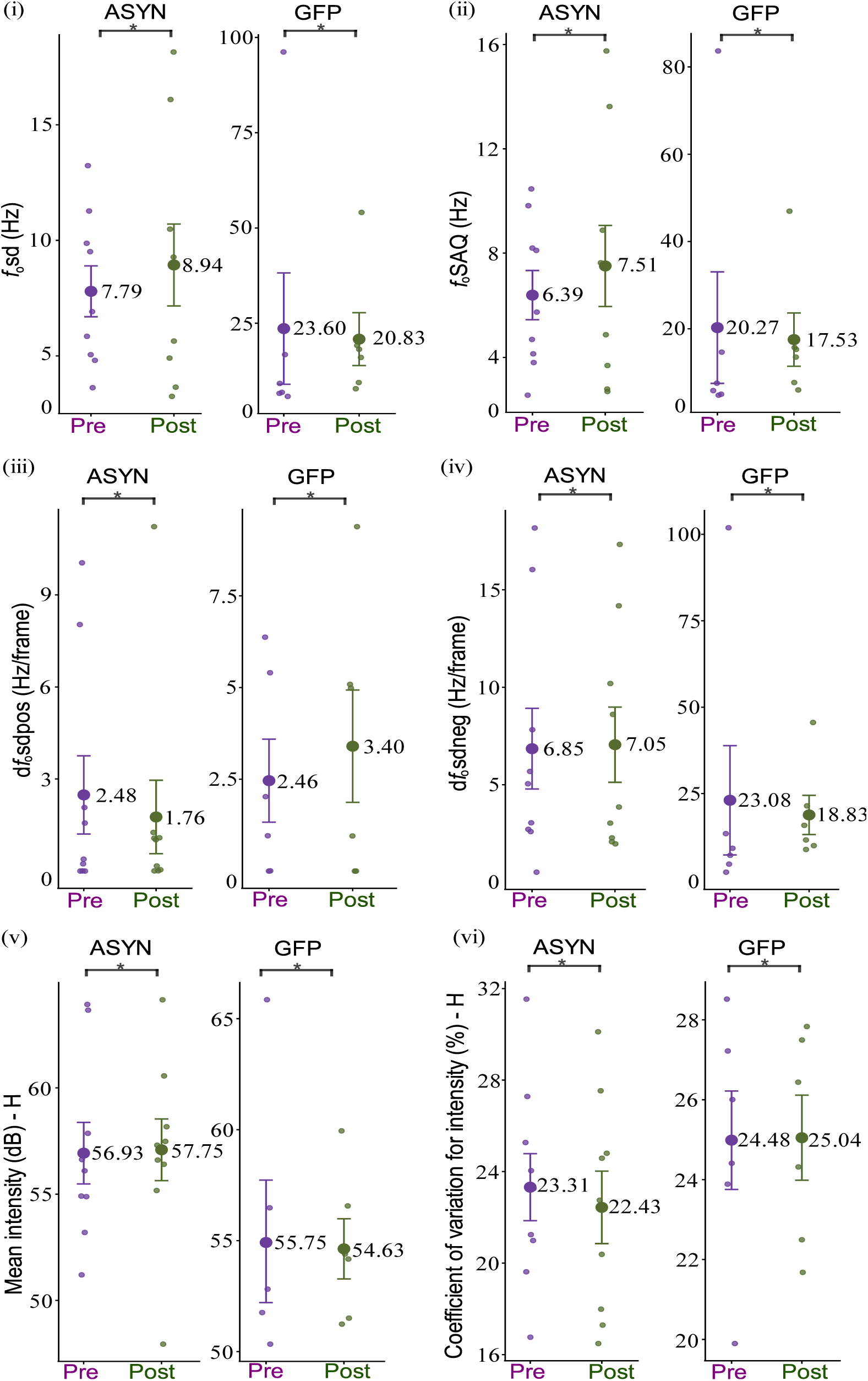
Representative *f*_o_ variability and intensity measures for harmonic syllables. Individual and group-level changes in acoustic-prosodic measures from pre- to post-intervention in the ASYN and GFP groups. Graphics show (i) *f*_o_sd, (ii) *f*_o_SAQ, (iii) d*f*_o_sdpos, (iv) d*f*_o_sdneg, (v) mean intensity (mint) for harmonic syllables, and (vi) coefficient of variation for intensity (cvint) for harmonic syllables. Small colored points indicate individual bird means at the pre- and post-virus injection timepoints. Large points represent group means, and error bars indicate ±1 standard error of the mean. * Indicate statistically significant differences (p < 0.05). Data are shown separately for the ASYN and GFP groups.

## Supplemental Method

### Statistical Power Analysis (Pre-Post Comparisons)

For paired pre-post comparisons, statistical power was estimated using a sensitivity-based approach for the Wilcoxon signed-rank tests. Effect sizes (*r*) derived from standardized Z statistics were used to approximate expected statistical power under the observed sample sizes. Power values were computed for each parameter at the level of paired bird-wise means, reflecting the probability of detecting the observed within-subject effects given the sample size and effect magnitude. As with between-group analyses, these estimates are reported as descriptive indicators of analytical sensitivity rather than inferential outcomes.

